# A holobiont view of island biogeography: unraveling patterns driving the nascent diversification of a Hawaiian spider and its microbial associates

**DOI:** 10.1101/2020.12.07.414961

**Authors:** Ellie E. Armstrong, Benoît Perez-Lamarque, Ke Bi, Cerise Chen, Leontine E. Becking, Jun Ying Lim, Tyler Linderoth, Henrik Krehenwinkel, Rosemary Gillespie

**Affiliations:** Department of Biology, Stanford University, Stanford, CA, USA; Institut de Biologie de l’ENS (IBENS), Département de biologie, École normale supérieure, CNRS, INSERM, Université PSL, Paris, France; Institut de Systématique, Évolution, Biodiversité (ISYEB), Muséum national d’Histoire naturelle, CNRS, Sorbonne Université, EPHE, UA, Paris, France; Computational Genomics Resource Laboratory, California Institute for Quantitative Biosciences, University of California, Berkeley, CA, USA 94720; Museum of Vertebrate Zoology, University of California, Berkeley, CA, USA 94720; Ancestry, 153 Townsend St., Ste. 800 San Francisco, CA, USA 94107; Department of Environmental Science, Policy and Management, University of California, Berkeley, CA, USA; Long Marine Laboratory, University of California, Santa Cruz, CA, USA; Marine Animal Ecology Group, Wageningen University & Research, Wageningen, The Netherlands; Wageningen Marine Research, Den Helder, The Netherlands; School of Biological Sciences, Nanyang Technological University, 60 Nanyang Drive, Singapore 637551; Department of Genetics, University of Cambridge, UK; Department of Biogeography, Trier University, Trier, Germany

**Keywords:** Host-associated microbes, endosymbiont, speciation, population structure, adaptive radiation, Ariamnes, Hawaiian Islands

## Abstract

The diversification of a host lineage can be influenced by both the external environment and its assemblage of microbes. Here, we use a young lineage of spiders, distributed along a chronologically arranged series of volcanic mountains, to determine the parallels between the evolutionary histories of the host spiders and their associated microbial communities, together forming the “holobiont”. Using the stick spider *Ariamnes waikula* (Araneae, Theridiidae) on the island of Hawaiʻi, and outgroup taxa on older islands, we tested whether each component of the holobiont (the spider hosts, the intracellular endosymbionts, and the gut microbial communities) showed correlated signatures of diversity due to sequential colonization from older to younger volcanoes. In order to investigate this, we generated ddRAD data for the host spiders and 16S rRNA gene amplicon data from their microbiota. We expected sequential colonizations to result in a (phylo)genetic structuring of the host spiders and in a diversity gradient in microbial communities. Results showed that the host *A. waikula* is indeed structured by geographic isolation, suggesting sequential colonization from older to younger volcanoes. Similarly, the endosymbiont communities were markedly different between *Ariamnes* species on different islands, but more homogeneous among *A. waikula* populations on the island of Hawaiʻi. Conversely, the gut microbiota was largely conserved across all populations and species, which we suspect are generally environmentally derived. Our results highlight that the different components of the holobiont have responded in distinct ways to the dynamic environment of the volcanic archipelago, showing the necessity of understanding the interplay between components to better characterize holobiont evolution.

## Introduction

Patterns of biodiversity are influenced by both ecological and evolutionary processes operating within the dynamic context of a community (Weber et al. 2017). The external environment can serve to isolate populations for various periods, and select for traits that influence their evolutionary trajectory. At the same time, a given organism also represents a community itself by hosting a diverse array of microbial species, many of which perform essential functions for their host (Koskella & Vos, 2015). The importance of microbial communities for promoting the isolation of their hosts (Bordenstein et al. 2001) and facilitating their adaptation to novel ecological niches (O’Connor et al. 2014) has been increasingly recognized. A species’ response to dynamic changes in the environment can frequently be modulated by the “holobiont” of host and microbial associates (Margulis & Fester 1991, Moran et al. 2019). In some cases, this holobiont may even act as a unit of selection (Guerrero et al. 2013). Therefore, understanding whether host organisms and their associated microbial communities respond in a similar manner to external forces, is essential for identifying potential drivers of evolution (McFall-Ngai et al. 2013).

Among arthropods, associated microbial communities are often highly diverse assemblages, accounting for an extensive range of interactions with their host (Engel & Moran 2013). Many arthropods host different microbial communities in different body parts that occupy various niches such as the gut microbiota or intracellular endosymbionts (Hansen & Moran 2014). The composition of the gut microbiota is often determined by complex interactions of environment, diet, developmental stage, and host evolutionary history (Yun et al. 2014), contributing to various functions for the host ranging from nutrition to protection against pathogens (Engel & Moran 2013, Moran et al. 2019, Hammer et al. 2019). However, for some arthropod taxa, recent work also suggests that a large proportion of the gut microbiota is purely environmentally derived, highly transient, and does not always have apparent functional relevance (Hammer et al. 2017; Kennedy et al. 2020). In contrast, a functional reliance of the host on its microbial communities could result in more stable and predictable gut microbial communities, that may be conserved and co-evolve with their host (Engel & Moran 2013, Kwong et al. 2017).

In contrast to the gut microbiota, most endosymbionts are vertically-transmitted intracellular bacteria. They can comprise tightly coevolved taxa, supplying their host with essential nutrients, such as bacteria of the genus *Buchnera* in aphids (Koga et al. 2003). Many other endosymbionts manipulate the reproduction of their host, such as species in the genera *Wolbachia*, *Rickettsia*, *Rickettsiella,* and *Cardinium* (Duron et al. 2017; Hoy & Jeyaprakash 2005; Vanthournout & Hendrickx 2015; White et al. 2020; Zhang et al. 2017). These taxa can promote cytoplasmic incompatibilities between hosts, enhancing genetic isolation (Shropshire et al. 2020). Some endosymbionts have also been associated with behavioral tendencies for long-distance dispersal (Goodacre et al. 2006), translating into higher gene flow and suppressing divergence between lineages otherwise isolated by distance.

Considering this background, a key point of interest is dissecting to what extent the nature of the host-microbe relationship affects the assembly and composition of host-associated microbial communities as the host adapts and evolves in its environment. Here, we examine the assembly and composition of different microbial communities within an evolving host by focusing on a lineage of spiders that shows recent divergence between populations on the youngest island of Hawaiʻi (Gillespie et al. 2018).

The Hawaiian archipelago is comprised of a chain of volcanic islands that increase in age from southeast to northwest, providing an ideal system for tracking the interplay between a host and the different components of its microbial community within an isolated setting (Shaw & Gillespie 2016). Even on the youngest island of Hawaiʻi, the volcanoes are arranged chronologically, from approximately 430,000 years old (Kohala), to the active flows of Kīlauea, with population structure of local flora and fauna generally shaped by progressive colonization of the newly emerged volcanoes (e.g. Simon 1987; Wagner & Funk, 1995; Blankers et al. 2018; Eldon et al. 2019; Goodman et al. 2019). Stick-spiders in the genus *Ariamnes* (Theridiidae) have diversified rapidly across the landscapes in the Hawaiian archipelago (Gillespie & Rivera 2007) and exhibit repeated diversification into ecomorphs adapted to specific microhabitats such as the “gold” ecomorph that is are entirely cryptic on the underside of leaves (Gillespie & Rivera 2007; Gillespie *et al*. 2018). However, their diet is conserved and they are specialized consumers of other spiders (Kennedy et al. 2018). Compared to their mainland counterparts in which dispersal by ballooning has been well documented (e.g. Larrivee & Buddle 2011), Hawaiian *Ariamnes* (as with other Hawaiian spiders) show little propensity for dispersal; such a loss of dispersal, and associated high levels of local endemism, are a common feature of organisms on remote islands (Gillespie et al 2012).

The current study focuses on a single species of the Hawaiian *Ariamnes* (*A. waikula*), endemic to the youngest island of Hawaiʻi. We aim to determine whether this highly specialized spider lineage shows parallel signatures of recurrent colonization events across volcanoes within the island in the same way as its microbial associates. We hypothesize that the population structure of *A. waikula* reflects a stepping stone colonization from older to younger volcanoes, and that populations from geologically older sites will show higher differentiation and higher within population diversity compared to younger sites, as they generate novel diversity subsequent to the founder event(s) (Roderick et al. 2012). For the microbial communities within the spider hosts, patterns of composition may or may not reflect the patterns of host differentiation depending on the nature of the host/microbe relationship. First, we predict that microbial associates that are vertically transmitted, including the intracellular endosymbionts, (i) will show patterns of community composition that closely mirror the population structure of the host (i.e. strong correlation between the host genetic structure and the microbial beta diversities) and (ii) will present lower microbial alpha diversities at the younger sites due to stochastic losses of some microbial symbionts during colonization events (Minard et al. 2015). Second, if microbial associates are mainly environmentally acquired and filtered by phylogenetically-conserved host traits (Moran & Sloan 2015), we can expect a correlation between the host genetic structure and the microbial beta diversities (Mazel et al. 2018) and we do not expect any reduction of the microbial alpha diversities at the younger sites. Third, if host-associated microbial communities are passively assembled without any filtering by the host, we expect no particular patterns in terms of microbial alpha diversity or relating host genetic structure and the microbial beta diversities. In the latter case, microbial beta diversities might rather be linked to geographical distances or other environmental factors.

To test these predictions, we examined the population genetic structure of *A. waikula* on Hawaiʻi Island, along with several outgroup species from other islands, using genome-wide single nucleotide polymorphism (SNP) data generated with double digest RAD sequencing (ddRAD). We then investigated how different components of their microbiota have changed as the spiders colonized new locations. To do so, we compared the community structure of microbial populations to that of their host individual using 16S rRNA gene amplicon sequencing, capturing the diversity of both the endosymbionts and the gut microbiota. Finally, we complemented these analyses with simulations to assess the robustness of our findings.

## Materials and Methods

### Sampling

We sampled *Ariamnes* across Hawaiʻi Island, focusing on individuals of *A. waikula* (gold ecomorph), from 6 populations. In addition, we included 2 individuals of the related *A. hiwa* (brown ecomorph) and 32 gold ecomorph individuals from two other species: *A. melekalikimaka* on West Maui and *A. n. sp.* Molokaʻi (Gillespie et al. 2018). Using *A. hiwa*, *A. melekalikimaka*, and *A. n. sp.* Molokaʻi we were able to confirm monophyly of the Hawaiʻi Island clade and compare the diversity of the microbial communities of other species between and within islands. Sampling sites were all located within the wet forest (rainfall 2500-3000mm) and between elevations of 1000m and 1300m, with the exception of the “Saddle” population which was slightly higher (Table S1). Our sites were all chosen to standardize variables (temperature, precipitation, elevation) to the extent possible. Individuals were collected by hand and immediately preserved in 90% EtOH. We collected a total of 133 individuals for sequencing (Supplementary Tables S1 & S2). Only adults were collected for this study, to decrease the likelihood of capturing differences driven by age.

**Table 1:**
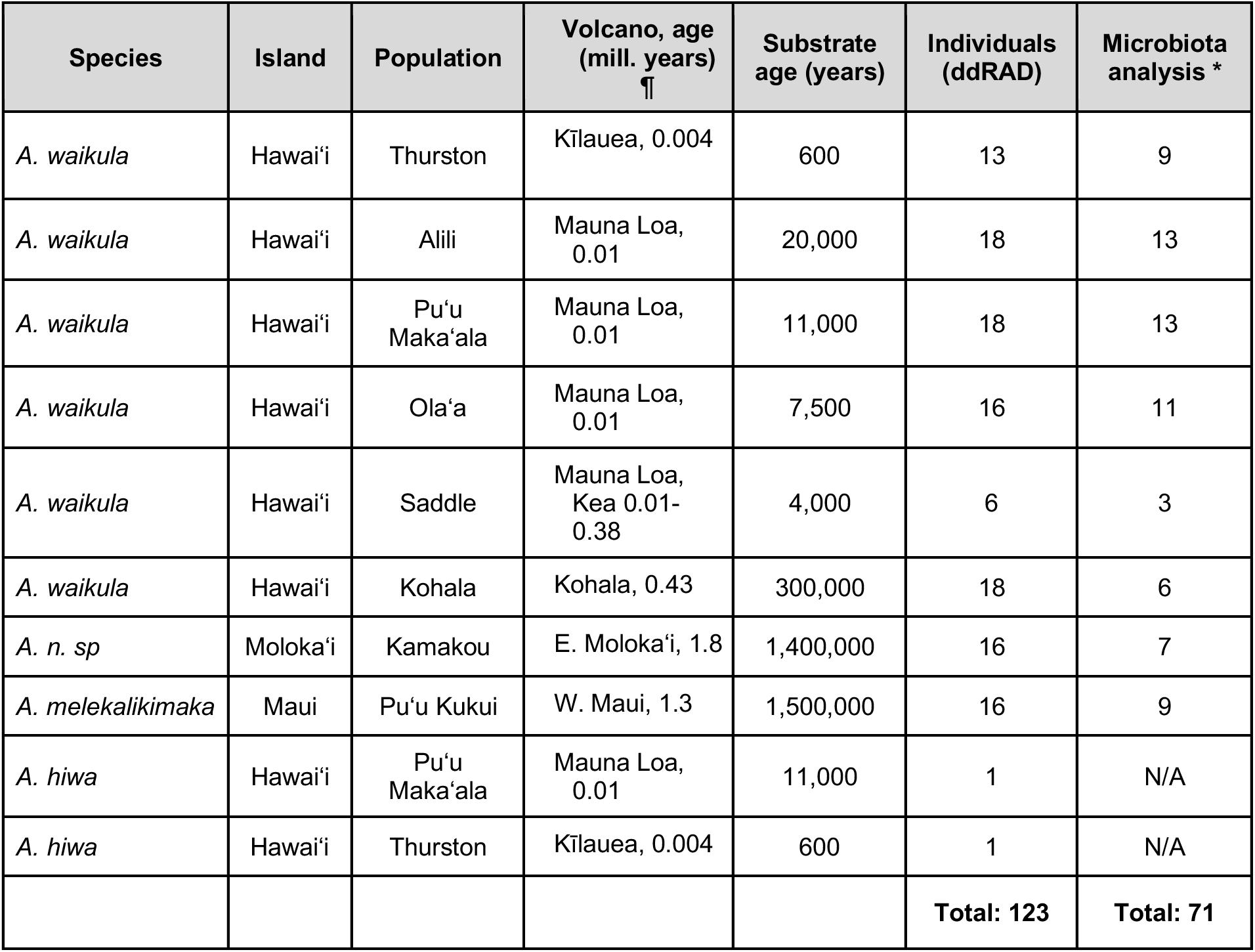
Individual, population, and sampling locality information for *Ariamnes* spiders in this study. Volcano ages are given based on K-Ar dating estimates of the maximum time available for occupancy (Langenheim & Clague 1987). Substrate ages (in years) are approximate and based on geologic estimates of the youngest lava flow making up each sampling locality. * Samples with sufficient coverage for microbiota analysis. ¶ Approximate ages and estimates of time available for colonization (Carson & Clague 1995). For full details see Supplementary Tables S1 and S2.

### Ariamnes - ddRAD library preparation and sequencing

To examine the population structure of the spider host, we used ddRAD to obtain reduced representation genome-wide SNP data. Genomic DNA was extracted from spider legs with several modifications to the Qiagen DNeasy kit protocol. Legs were first removed from each specimen using sterile tweezers so that the abdomen remained intact for the microbial DNA analysis. DNA was then extracted by placing the tissue in Proteinase K and lysis buffer and grinding them with a sterile pestle to break up the exoskeleton. We then added 4uL RNase A (100mg/ml) and incubated the extractions for two minutes at room temperature. Tubes with tissue and extraction solution were then placed overnight in a heat block at 56°C. The remainder of the extraction protocol was performed following the manufacturer’s instructions. We built ddRAD libraries following an adapted protocol of Peterson *et al*. (2012) (Saarman & Pogson, 2015; see Maas et al. 2018 for protocol optimization steps). Briefly, we started the ddRAD protocol with a total of 100 nanograms of DNA per sample. The DNA was digested using SphI-HF (rare-cutting) and MlucI (frequent-cutting) restriction enzymes. We assessed fragmentation with a Bioanalyzer High Sensitivity chip (Agilent). We multiplexed 15-20 individuals per library for a total of eight ddRAD libraries, using oligo sequences of 42bp including a barcode of 5bp (Peterson *et al*. 2012). We used a Sage Science Pippen Prep to size select 451-551bp (including internal adapters) fragments, and confirmed the sizes using a Bioanalyzer. Ten indexing polymerase chain reaction (PCR) cycle were run on each library to enrich for double-digested fragments and to incorporate a unique external index for each library pool. The eight libraries were sequenced using 100bp paired-end sequencing on one Illumina HiSeq2500 lane at the Vincent J. Coates Genomic Sequencing Facility at UC Berkeley.

### Ariamnes - ddRAD data filtering and processing

We used a custom perl script invoking a variety of external programs to filter and process the ddRAD data (RADTOOLKITv0.13.10; https://github.com/CGRL-QB3-UCBerkeley/RAD). Briefly, raw fastq reads were first de-multiplexed based on the sequence composition of internal barcodes with tolerance of one mismatch. De-multiplexed reads were removed if the expected cutting site was not found at the beginning of the 5’-end of the sequences. The reads were then filtered using cutadapt (Martin 2011) and Trimmomatic using default parameters (Bolger et al. 2014) to trim off Illumina adapter contaminations and low-quality reads. The resulting cleaned forward reads of each individual were first clustered using cd-hit with 97% as the clustering threshold (Fu et al. 2012; Li & Godzik 2006), keeping only those clusters with at least two reads. In each cluster, we pulled out the corresponding reverse reads based on the identifiers. Both forward and reverse clusters were kept only if the corresponding reverse reads also formed one cluster. If reverse reads were grouped into more than one cluster, then only the forward read cluster was kept. For each paired cluster, the representative sequences for each forward and reverse cluster determined by cd-hit were retained. We then merged the forward and reverse sequences using FLASH (Magoč & Salzberg 2011). If they could not be merged then they were joined by placing “N”s between the two sequences. The resulting loci were then masked for putative repetitive and low-complexity elements, and short repeats using RepeatMasker (Smit et al. 2004) with “spider” as the database. After masking, we eliminated loci if more than 60% of the nucleotides were Ns. The resulting ddRAD loci from each individual were combined and clustered for all individuals. Contigs that were at least 40 nucleotides in length and shared by at least 60% of all the individuals served as a reference. Cleaned sequencing reads from each individual were then aligned to the reference using Novoalign (http://www.novocraft.com), discarding any ambiguously mapped reads. We used Picard (http://picard.sourceforge.net) to add read groups and GATK (McKenna et al. 2010) to perform realignment around indels in BAM format generated by SAMtools (Li et al. 2009). We then used SAMtools/BCFtools (Li et al. 2009) to generate data quality control information in VCF format. These data were then further filtered using a custom perl script, SNPcleaner (Bi et al. 2013). We filtered out any loci with more than two called alleles. We masked sites within 10 bp upstream and downstream of an indel. We discarded sites with a total depth outside of the genome-wide 1st and 99th percentile. To avoid excessive heterogeneity in sample representation among sites, we also removed alignments if more than 40% of the samples had less than 3X coverage.

Since most of the individuals were expected to have low coverage (between 5-15x) we used ANGSD (Korneliussen et al. 2014) to calculate genotype likelihoods and genotypes for the analyses. This tool was specifically developed for population and evolutionary genomic analyses of low coverage data. We used genotype likelihoods whenever the downstream tools allowed us to, randomness in allele sampling as well as sequencing or mapping errors lead to uncertainty in genotype calling (Crawford & Lazzaro 2012)).

### Ariamnes - Phylogenetic Analyses

We investigated the phylogenetic relationships between populations of *A. waikula* and the other *Ariamnes* species to better understand colonization patterns. To do so, we used the Stacks pipelines (Catchen et al. 2013) to group reads into homologous loci across all individuals and extract phylogenetically informative sites (i.e. fixed within individuals but variable between individuals). This yielded an alignment of 58,899 sites with which we performed phylogenetic reconstruction using IQ-TREE (Nguyen et al. 2015) by combining model selection from ModelFinder Plus (Kalyaanamoorthy et al. 2017) and assessing branch supports with 1,000 ultrafast bootstrap (Hoang et al. 2018). Finally, we rooted the tree using the *A. hiwa* individuals and calibrated it with r8s (Sanderson 2003) without specifying any absolute dating (*i.e.* setting the root age to 1).

### A. waikula *-* Population Genetic Analyses

We first explored *A. waikula* population structure using Principal Components Analysis (PCA). PCA, as opposed to structure analyses which force data into a specific number of clusters, is an unsupervised method which attempts to find orthogonal directions of maximum variance through a cloud of data. Therefore, it is a useful way to explore data structure prior to using supervised methods for inferring population structure. We used the previously generated genotype likelihoods to calculate genotype posterior probabilities under an allele frequvvency prior (-doPost 1) in binary format (-doGeno 32) for use with ngsCovar (part of the ngsTools package; (Fumagalli et al. 2014). We used ngsCovar to calculate a genetic covariance matrix among individuals from the genotype probabilities at sites having a minor allele frequency of 5% to avoid noise from very rare alleles (which could be due to sequencing errors). Comparisons between the first three principal component axes were then plotted using R (R Core Team, 2020). Pairwise F_ST_ values were calculated in ANGSD from allele frequency likelihoods (-doSaf 1) and the respective pair’s genome-wide, unfolded, joint site frequency spectrum (SFS) as a prior for jointly observing any combination of allele frequencies between the two populations. We used unfolded allele frequencies by supplying the reference sequences as a pseudo-ancestral sequence since ANGSDv0.930 is only able to accurately estimate F_ST_ using unfolded data. In order to visualize F_ST_ distances, we performed multidimensional scaling (to 2 dimensions) with R (R Core Team, 2020; using the *cmdscale* function). Last, we estimated pairwise genetic distances calculated from genotype likelihoods using the software ngsDist (Viera et al. 2015) and balanced minimum evolution using FastME (Lefort et al. 2015). To do so, we first calculated genotype likelihoods using ANGSD (see above), which were used to construct a pairwise distance matrix between all individuals using ngsDist.

A crucial factor affecting population structure is gene flow between populations. In order to determine levels of connectivity between populations of *A. waikula*, we used two different approaches: ngsAdmix (Skotte *et al*. 2013) and EEMS (Petkova et al. 2016). ngsAdmix is a genotype likelihood based-tool for estimating individual admixture proportions, while EEMS uses genotype calls to infer effective migration surfaces. We inferred ancestry proportions indicative of admixture for different values of K (number of ancestral populations; ranging from two to six, which is the number of populations sampled) using R for visualization. We then generated genotype calls for the EEMS analysis using ANGSD with the following flags: ‘-doMaf2’, ‘-doGeno 2’, ‘-doPost 1’, ‘-doSaf 1’, ‘-fold 1’, a SNP p-value cut-off of 1e-6, and a posterior probability for calling genotypes of 0.75. We only considered sites with a minimum depth of 3. We converted the output into the adegenet (R-package) input format using PopGenTools (https://github.com/CGRL-QB3-UCBerkeley/PopGenTools) to calculate genetic distances between all individuals. EEMS also requires a geographic map of the area in question, so we used the program to divide Hawaiʻi Island into grids of 10km (Supplementary Fig. S1) and subsequently added the samples to the grid using one GPS coordinate per population (Supplementary Table S1). Using a stepping stone model, we calculated migration rates between the demes. Convergence of the MCMC runs was assessed by plotting and inspecting the traces by eye (Supplementary Fig. S2). We performed 10 independent runs for each of the analyses.

Next, we calculated population genetic statistics such as nucleotide diversity (Pi), Watterson’s Theta, and Tajima’s D. To do so, we generated a folded SFS for each population using ANGSD and realSFS (part of the ANGSD package). We then ran ANGSD *-doTheta 1* and *-pest* (which provides the genome-wide SFS prior to ANGSD) and the respective output formats were converted to bed format using *thetaStat make_bed*. Subsequently, we calculated the per population statistics using *thetaStat do_stat*. The average, min, and max Tajima’s D were then calculated and we further extracted the genome-wide average Watterson’s theta and Pi values. While Watterson’s theta and Pi are both measures of genetic diversity, Watterson’s theta is based only on the number of segregating sties and so is sensitive to novel, rare diversity, whereas Pi is most influenced by sites with alleles at intermediate frequency more typical ancestral diversity. We measured both Pi and theta, as well as Tajima’s D to gain a wholistic understanding of the genetic signatures of colonization events and possible subsequent expansions. We then also used linear models in R to test whether genetic diversity (Pi or Watterson’s theta) was positively related to the age of a given volcano.

### Microbial Communities

To characterize the microbial community within the *A. waikula* hosts, a subset of 71 individuals from the eight populations on Hawaiʻi and two additional islands (Table 1; Supplementary Tables S1 & S2) were selected for analysis. We focused on the mid and hindgut, both located in the spider’s opisthosoma. The preservation in ethanol led to considerable shrinkage of the opisthosoma and thus did not allow us to separately dissect out the gut. Instead, we used the whole opisthosoma to extract DNA, as described in Kennedy et al. (2020). Specimens which did not have the opisthosoma intact were not used. The digestive tract comprises the majority of the opisthosoma’s cavity. In addition, it contains silk glands, the heart, lungs, and gonads. The opisthosoma was removed with a sterile razor blade and then washed in ethanol to remove external bacteria (Hammer et al. 2015). Besides the intracellular endosymbionts, we considered the remaining bacteria to be representative of the “gut microbiota”, even if it technically consists of the “opisthosoma microbiota”, but previous studies have shown that the gut microbiota dominates in the opisthosoma (Sheffer et al. 2020, Kennedy et al. 2020). The tissue was then transferred into lysis buffer and finely ground with a sterile pestle. DNA was extracted using the Gentra Puregene Tissue Kit (Qiagen, Hilden, Germany) according to the manufacturer’s protocol. Spider abdominal tissue can contain PCR inhibitors (Schrader et al. 2012), thus we cleaned the DNA extract with 0.9X AmPure Beads XP.

We next amplified a ∼300 bp fragment of the V1-V2 region of the bacterial 16S rRNA using the Qiagen Multiplex PCR kit according to the manufacturer’s protocols and using the primer pair MS-27F (AGAGTTTGATCCTGGCTCAG) and MS-338R (TGCTGCCTCCCGTAGGAGT) (Gibson et al. 2014). PCRs were run with 20ng of template DNA and 30 cycles at an annealing temperature of 55°C. PCR products were separated from leftover primer by 1X AmPure Beads XP. A six-cycle indexing PCR was performed on the cleaned products, adding dual indexes to every sample using the Qiagen Multiplex PCR kit. Indexing was performed according to (Lange et al. 2014). The dual indexed libraries were isolated from leftover primer as described above, quantified using a Qubit fluorometer, and pooled in equal amounts into a single tube. The library was sequenced on an Illumina MiSeq using V3 chemistry and 300 bp paired reads. In order to discard contaminants from our final dataset, we also performed blank extraction controls and negative PCR controls (without DNA template), which were sequenced along the other samples.

We used the 16S profiling analysis pipeline for Illumina paired-end sequences of the Brazilian Microbiome Project (Pylro et al. 2014), including QIIME 1.8.0 (Caporaso et al. 2010) and USEARCH 11 (Edgar 2013). We modified some steps of these pipelines, using our own Bash and R scripts (R Core Team, 2020; see Data Accessibility section). The QIIME script join_paired_ends.py was used to merge paired reads. The fastq_filter command in USEARCH was used for quality filtering the assemblies (below a base calling error probability of 0.5, using an average Q-score for each read). We used the Stream EDitor in UNIX to remove PCR primers from all assembled sequences. Sequences were de-replicated using USEARCH, removing all singletons. OTUs were generated at a similarity cutoff of 3% or 0% (0 radius OTUs, or “Z-OTU”) from the de-replicated sequences and chimera removed de novo using USEARCH. The following analyses were thus independently applied on the two distinct sets of OTUs. We assigned taxonomy to the resulting OTUs using the assign_taxonomy.py script based on the Greengenes database (http://greengenes.secondgenome.com). We removed sequences corresponding to OTUs found in high prevalence and abundance in the different negative controls from all samples, as these could represent contaminants (Hornung et al. 2019).

Spiders are known to carry various endosymbiotic bacteria (Goodacre et al. 2006, Vanthournout et al. 2015, White et al. 2020). These can be vastly overrepresented in microbial analyses and thus may completely dominate the microbial community structure. Since we did not extract the gut from individuals, endosymbionts from outside of the gut could be particularly prevalent in our analysis. We thus separated known endosymbionts in our samples (*i.e. Rickettsia, Rickettsiella, Wolbachia,* and *Canditatus Cardinium*) from the remaining bacterial sequences (referred to as the “gut microbiota”), resulting in two OTU sequence files. Both of these datasets were used for the following analyses separately.

OTU tables were prepared by mapping sequences back to the filtered OTU sequence files using USEARCH. The OTU tables were rarefied to an even coverage (from 400 to 8,000 reads with 20 replications per rarefaction depth) using the multiple_rarefactions.py script in QIIME. Given the rarefaction curves (Supplementary Fig. S3), we chose rarefied depths at 3,200, 110, and 3,000 reads for the whole microbiota, the endosymbionts, and the gut microbiota respectively and replicated 20 times this rarefaction; these rarefied OTU tables were used in all the following analyses. Rarefying at 110 reads for the endosymbionts enabled us to keep all the samples for the following analyses, but using a higher threshold did not qualitatively affect our results (not shown) given that most samples were dominated by a single endosymbiotic OTU (see Results). We plotted the relative abundances of different microbial taxa at the genus and order levels for the different *Ariamnes* populations.

We then investigated whether microbial associates showed similar diversity patterns compared to their *Ariamnes* hosts along the chronosequence. Chao1 indexes of alpha diversity were calculated from the rarefied OTU tables using QIIME (alpha_diversity.py), and to evaluate the presence of a bottleneck in microbial diversity, linear mixed models were used to test for an effect of the volcano age and the genetic diversity of *Ariamnes* host populations (Pi and Watterson’s theta) on the bacterial alpha diversity, with sampling site as a random effect. The different variables were log-transformed prior to analyses, and homoscedasticity and normality of the model residuals were verified.

To measure microbiota differentiation across *Ariamnes* species and populations, we computed beta diversity between microbial communities of each individual using QIIME (beta_diversity.py with Bray-Curtis dissimilarities). Beta diversity of microbial populations was visualized with a Principal Coordinate Analysis (PCoA) and as dendrograms using a neighbor joining reconstruction with the R-package ape (Paradis et al., 2004). To test whether individuals from the same population tend to host similar bacterial communities across the different islands (*i.e.* different *Ariamnes* species) or within Hawaiʻi Island only (*A. waikula*), we performed a Permutational analysis of variance (PERMANOVA; *adonis* function, vegan R-package) on the beta diversity matrices with 10,000 permutations, after having verified the homogeneity of the variances (*betadispers* function). Next, we analyzed whether the microbiota of the *Ariamnes* holobiont mirror the host’s phylogeny by testing the correlations between microbial Bray-Curtis dissimilarities and host genetic distances (ngsDist distances) or between microbiota differentiation and host phylogenetic distances, using Mantel tests with 10,000 permutations (vegan R-package). These analyses were performed between populations of *A. waikula* in Hawaiʻi Island and were compared to the analyses performed between populations of different *Ariamnes* species (*i.e.* including *A. melekalikimaka* on West Maui and *A. n. sp.* Molokaʻi). Finally, we tested whether the microbiota differentiation could be explained by geography alone, by performing a Mantel test between the microbial Bray-Curtis dissimilarities and the geographical distances between sampling sites.

In addition, we replicated all diversity analyses of the microbial communities using phylogenetic diversity indexes: alpha diversities were measured using Faith’s phylogenetic diversity, and beta-diversities were measured using weighted UniFrac distances (Lozupone & Knight, 2005).

During all analyses of diversity, we also controlled for batch effects during the PCR steps by assessing the correlations between proximal samples in the PCR plates.

### Simulations

Our data for the microbial communities relied on 16S rRNA, raising the question as to whether the rate of evolution of this particular molecular marker was sufficient to capture changes at the very short timescales (<2 Myr) involved in the current study. The 16S rRNA gene of symbiotic bacteria is generally assumed to accumulate only 1-2% of sequence divergences per 50 Myr on average (Moran et al. 1993; Ochman et al. 1999). To investigate whether such a slow substitution rate is sufficient to detect microbial differentiation on <2Myr timescales we performed simulations. We directly modeled the evolution of the DNA sequences of transmitted OTUs along the *Ariamnes* phylogeny so as to simulate their evolution on a phylogenetic tree independent from the *Ariamnes* tree (following Perez-Lamarque & Morlon 2019). We considered DNA sequences of 300 bp, with 10% variable sites, and a substitution rate of 0.001 events per site per million years (such that two sequences would accumulate 1% divergence in 50 Myr on average). The *Ariamnes* microbiota were composed of 100 OTUs with a certain percentage of vertically transmitted OTUs (either 0, 10, 25, 50, 75, or 100%) and the remainder acquired from the environment. For each percentage of transmitted OTUs, we performed 30 simulations of mock microbiota using the function *simulate_alignment* from the R-package HOME (Perez-Lamarque & Morlon 2019). Then, we considered that each OTU had an abundance of 100 and we added a sampling process of 50% of the OTUs in each microbiota (representing the sampling hazards of a metabarcoding sequencing). Finally, we computed the Bray-Curtis beta diversities of the mock *Ariamnes* microbiota and tested the presence of microbial differentiation by performing a Mantel test with the host phylogenetic distances.

We then tested whether the presence of a bottleneck in the pool of available microbes along the island chronosequence could be detected in the alpha diversities of the *Ariamnes* microbiota. We considered 100 OTUs (each OTU having an abundance of 100) and simulated bottlenecks of the microbial OTUs between the old sites (Molokaʻi and Maui islands) and the younger Hawaiʻi Island. We considered different strengths of bottlenecks (from 0% to 50% of the microbial OTUs) and simulated mock microbiota for each *Ariamnes* by randomly picking 100 microbes from the local pools. We then tested the relationship between microbial alpha diversities measured using Chao1 index and the volcano age using linear mixed models. Note that here we only considered microbial bottlenecks as the stochastic losses of some microbial OTUs during island colonization, but the signal of the microbial bottlenecks can also be reinforced by the reduction of the genetic diversity within each microbial lineage at colonization events.

## Results

### Ariamnes - Phylogenetic Analyses

We obtained approximately 210 million paired-end reads from Illumina sequencing of the ddRAD libraries. After filtering for contaminants and low-quality reads, there were approximately 83 million remaining across the 123 demultiplexed samples. From these, we retained a total of 2,957,301 sites passing filters out of a possible 7,378,384 across all individuals that were used in downstream analyses and resulting in 123 individuals for population genetic analysis. After quality filtering, the average sequencing depth of samples was 12x (range of 3-42x across 123 samples).

Phylogenetic analysis confirms that the *A. melekalikimaka* (West Maui), *A. n. sp.* (Molokaʻi) and *A. waikula* (Hawaiʻi Island) are monophyletic groups (Fig. 1). Among *A. waikula* individuals, the population from Kohala (the oldest volcano on Hawaiʻi Island) is sister to the clade that contains all other individuals. Except for one individual from Saddle and two from Alili, these populations from Saddle (between Mauna Loa and Mauna Kea) and Alili form monophyletic clades, suggesting that there may be infrequent dispersal between some populations. The populations from Olaʻa, Puʻu Makaʻala and Thurston, all sites that are close together in the saddle between the volcanoes of Mauna Loa and Kilauea (MLKS), form one mixed clade.

**Figure 1:**
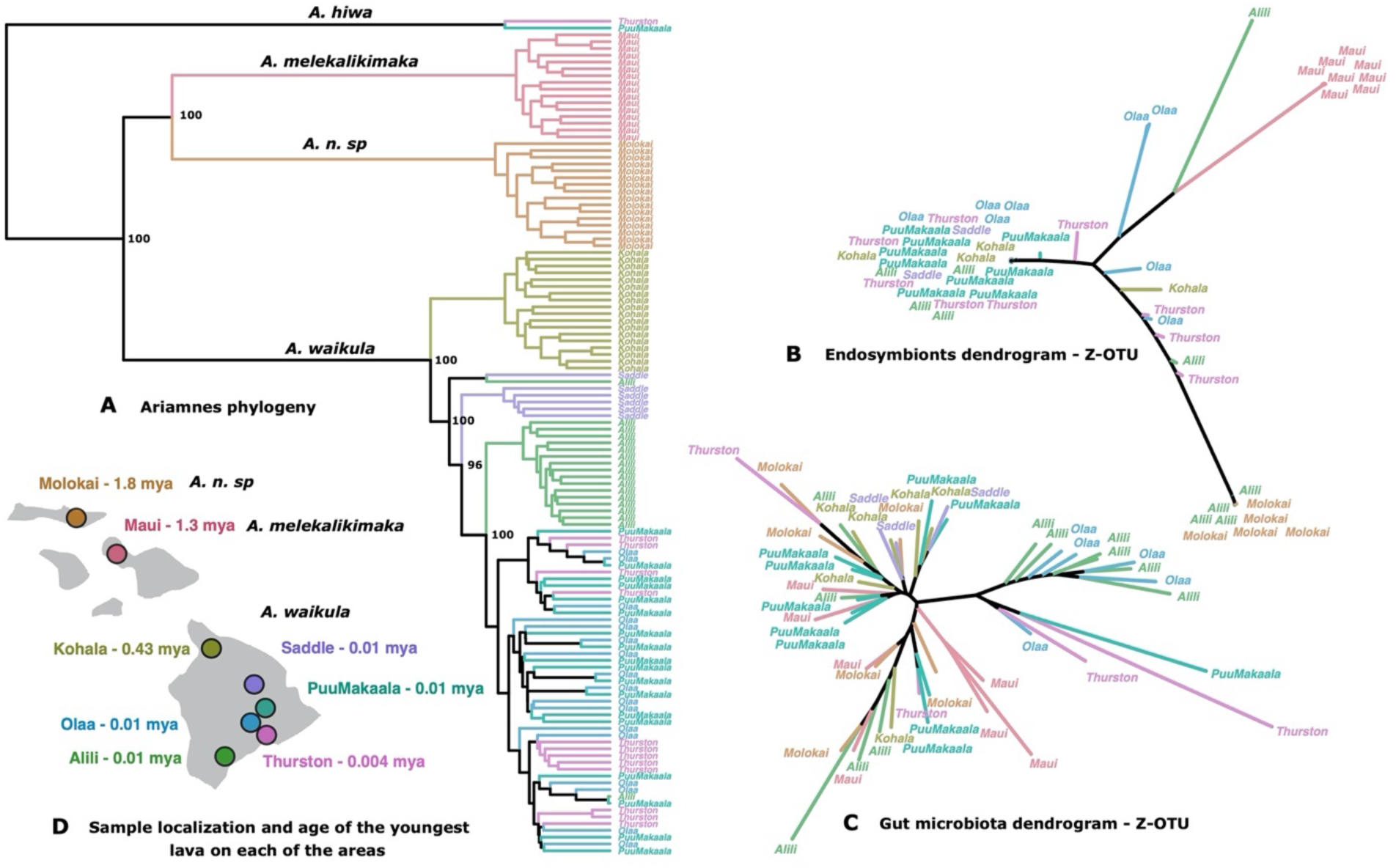
Host phylogenetic history partially recapitulates microbiota differentiation. (A) Phylogenetic tree of the *Ariamnes waikula* individuals across the island of Hawai’i obtained using IQ-TREE smoothed with r8s. Two specimens of *A. hiwa* (brown ecomorph) and 32 specimens of *A. melekalikimaka* (from Maui; gold ecomorph) and *A. n. sp.* (from Molokaʻi; gold ecomorph) were used as outgroup taxa. The bootstrap support values are indicated on the side of the main phylogenetic nodes. (B) Microbiota dendrograms reconstructed from the endosymbiont community and (C) the gut microbiota for the Z-OTU. (D) map of the sampled host populations and the corresponding volcano age on each of the areas.

### A. waikula - Population genetics

We analyzed population differentiation and structure for the full data set, and a subset including only *A. waikula* from Hawaiʻi Island. We did not include the two outgroup *A. hiwa* individuals. Divergence (F_ST_) was highest and similar between the different species on separate islands: 0.52-0.54 for *A. melekalikimaka* (Maui) compared to *A. waikula* (Hawaiʻi Island); 0.46-0.49 for *A. n. sp.* (Molokaʻi) compared to *A. waikula (*Hawai’i Island); 0.47 for *A. n. sp.* (Molokai) compared to *A. melekalikimaka* (Maui) (Table S3). These levels of divergence are close to those reported for COI data between *Ariamnes* populations on Maui compared to Hawaii Island (Roderick et al. 2012). *A. waikula* populations on Hawaiʻi Island were structured primarily according to locality, with Kohala, Alili, and Saddle being distinct from each other and from the MLKS sites (pairwise F_ST_ range 0.03-0.15; Puʻu Makaʻala, Thurston and Olaʻa; Fig. 1A).

Clustering analyses (PCA and ngsAdmix) of *A. waikula* broadly showed clustering of groups according to locality/volcano, except at MLKS sites (Fig. 2a, Supplementary Fig. S5). PC1 explained approximately 16% of the variation and separated Kohala and the younger sites, placing Saddle in the middle (Fig. 1A). Similarly, admixture analysis showed Kohala first splitting off at K=2. PC2 explained approximately 4% of the variation and primarily separated Alili from the MLKS sites, while PC3 (3%) separates Saddle, and to a lesser degree, Alili from the other populations (Supplementary Fig. S5). Admixture analyses show the same pattern, with individuals from Alili clustering separately at K=3 and Saddle at K=4. At K=6 we see slightly more similar ancestry between individuals from Thurston and Olaʻa, than either of these two site with Puʻu Makaʻala (Fig. 2B). For the ngsAdmix analyses, we found consistent results from the 50 independent runs for higher values of K for the Hawaiʻi only sampling (Supplementary Fig. S4). Lastly, the EEMS analyses with *A. waikula* individuals indicate potential gene flow between the MLKS populations: Puʻu Makaʻala, Thurston, and Olaʻa (Fig. 2C).

**Figure 2:**
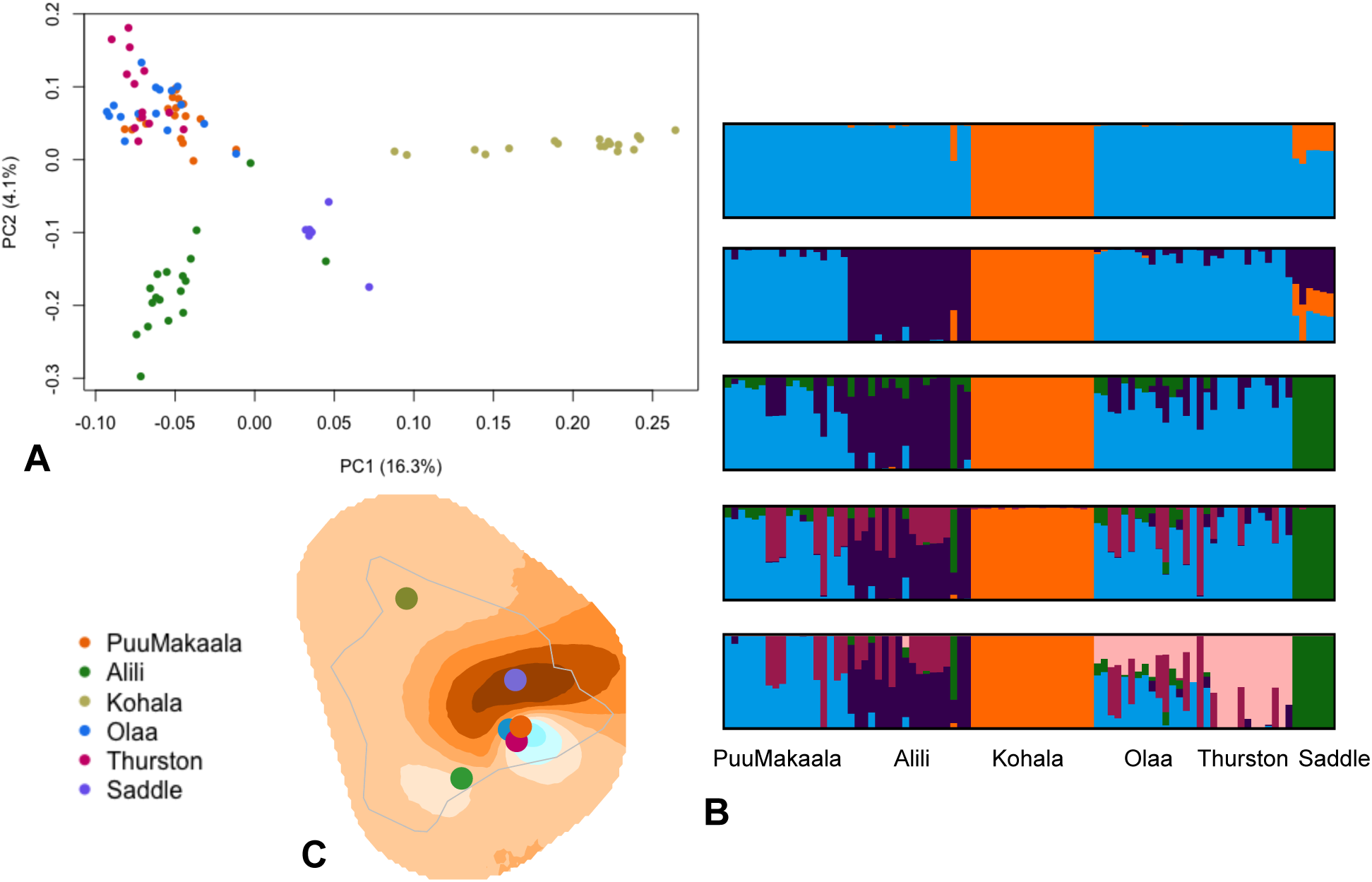
Population genetic analyses of *Ariamnes waikula*. A) ngsAdmix results for K=2 to K=8 for all *A. waikula* specimens. B) ngsAdmix results f or K=2 to K=6 (from top to bottom) for all *A. waikula* from Hawai’i Island only. C) EEMS analysis of all *A. waikula* specimens. Brown color indicates barriers for gene flow (the stronger the darker), and cyan indicates gene flow between populations (the stronger the darker).

Contrary to expectations, we found that Watterson’s theta was fairly even across localities on the youngest island of Hawai’i (Fig. 3). However, this correlation between Watterson’s theta and volcano age was not significant (F=0.023, p-value = 0.87) for *A. waikula* populations within Hawai’i (Figure 3) and no other correlations were found between theta and volcano age or for *Ariamnes* genetic diversity (Pi or Theta).

**Figure 3:**
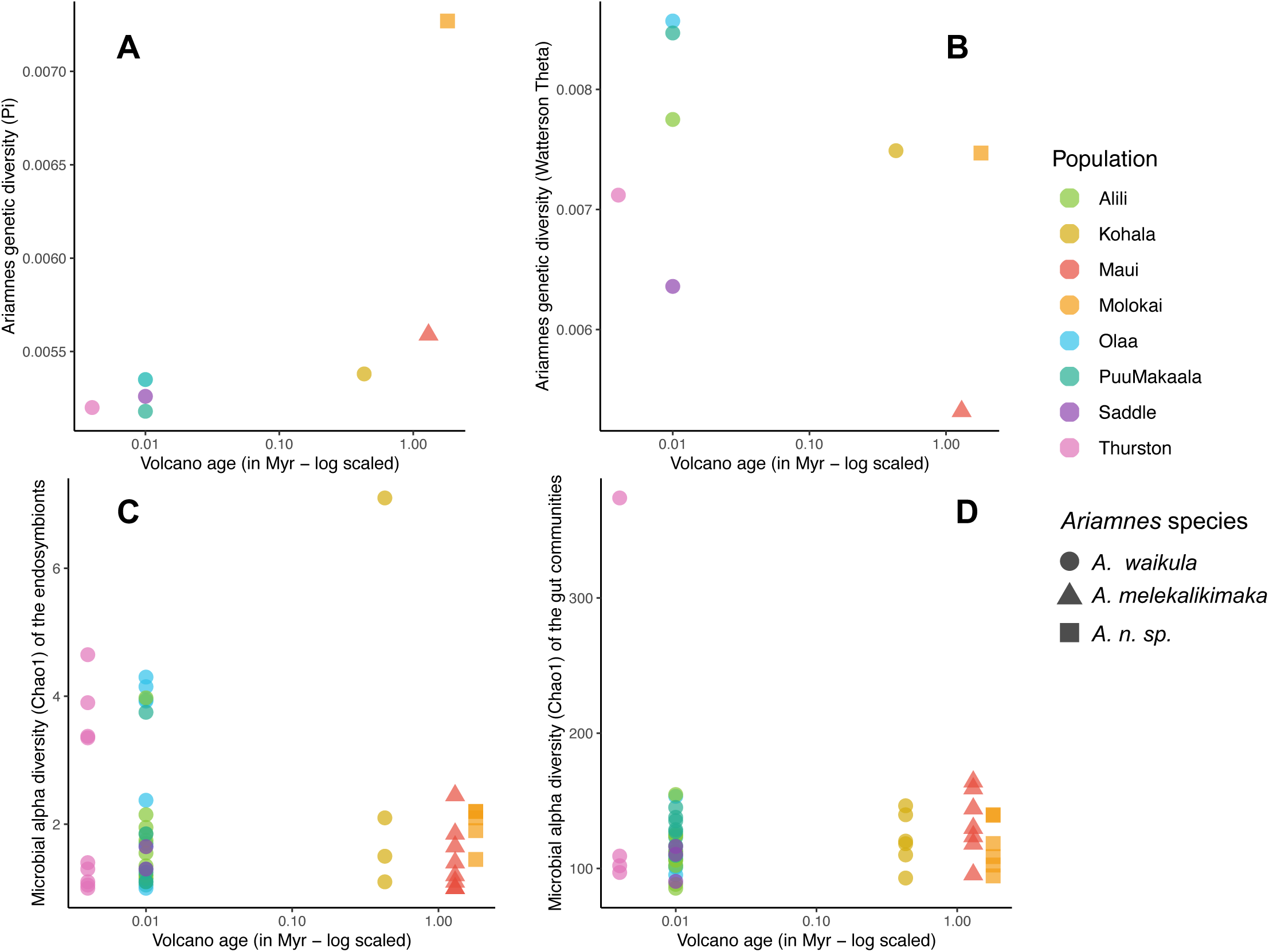
**The bottleneck in diversity along the volcano age tends to affect the *Ariamnes* hosts, but not their microbial associates** Note that Maui and Molokaʻi correspond to different *Ariamnes* species (*A. melekalikimaka* and *A. n. sp respectively*) and are only represented as a reference here: the following linear model testing the effect of volcano age on host (A-B) or microbial (C-D) diversities were only performed on the six *A. waikula* populations in Hawai’i. (A) *Ariamnes* genetic diversity (Pi) estimated per population as a function of the volcano age (in Myr). We found no significant relationship between *A. waikula* genetic diversity (Pi) and the volcano age (linear model: F=1.7, df=4, p-value=0.27), despite the observed trend, probably because of the small number of sampled populations in Hawai’i (6 only). (B) *Ariamnes* genetic diversity (Watterson’s theta) estimated per population as a function of the volcano ago (in Myr). We found no significant relationship between *A. waikula* genetic diversity (Watterson’s theta) and the volcano age (linear model: F=0.023, df=4, p-value=0.87). (C) Microbial alpha diversity (Chao1 index on Z-OTUs) estimated for the endosymbionts as a function of the volcano ago (in Myr). Each dot corresponds to the average of the 20 rarefactions performed for each individual. We found no significant relationship between endosymbionts alpha diversity and the volcano age (linear mixed model: t=1.10, p-value=0.24). (D) Microbial alpha diversity (Chao1 index on Z-OTUs) estimated for the gut community as a function of the volcano ago (in Myr). Each dot corresponds to the average of the 20 rarefactions performed for each individual. We found no significant relationship between gut microbial alpha diversity and the volcano age (linear mixed model: t=-0.08, p-value=0.94). The absence of a significant correlation was not influenced by the presence of an outlier sample in Thurston with very high microbial diversity.

We also investigated Tajima’s D, which reflects population size changes as a skew in the site frequency spectrum (Supplementary Fig. S6). We found that most sampling sites showed only marginally negative values consistent with population expansion (average: -1.0 to -0.06; Supplementary Table S4).

### Microbial Communities

We obtained a total of 4,932,236 bacterial reads from the 16S rRNA sequencing. After the quality filtering steps, demultiplexing, and removal of contaminants based on the negative controls (including the genera *Brachybacterium*, *Veillonella*, or *Brevibacterium*), we obtained a total of 1,315,469 sequences, which range from 5,000 sequences to 45,000 per individual *Ariamnes* sample. Due to the largely unexplored nature of the microbial communities of Hawaiian arthropods, we could not assign genus level taxonomy to the majority of microbial OTUs (only 241OTUs out of the 571 97% OTUs and 742 OTUs out of the 1,357 Z-OTUs), but were able to assign order for the majority of them (528 OTUs 97% and 1,290 Z-OTUs).

We did not find any significant differences in the alpha diversities of the whole microbiota among *Ariamnes* populations or species (Supplementary Table S5). Using linear mixed models, we also did not find a significant association of the microbial alpha diversity with volcano age or the genetic diversity of the host population (Fig. 3), even though the alpha diversities of the gut microbiota seemed to be slightly higher for host populations present on old volcanos (Fig. 3D). The results were similar for both Z-OTUs and 97% OTUs.

#### Intracellular endosymbiont community

When only selecting the endosymbionts, the community showed considerable variation between the different species of *Ariamnes* (Fig. 1B). *Ariamnes melekalikimaka* from West Maui mostly carry *Wolbachia*, while *Rickettsia* dominate the endosymbiont community of *A. n. sp.* Molokaʻi, and *Rickettsiella* dominates the populations of *A. waikula* on Hawaiʻi Island (Fig. 4A & Supplementary Fig. S7). This suggests that each spider population was colonized by a single endosymbiont strain (single OTU; Supplementary Fig. S8). The exception to this was several individuals of *A. waikula* spiders from Alili, Olaʻa, and Thurston that appear to have been simultaneously colonized by *Rickettsiella* and *Rickettsia.* We observed a large variability in the relative abundance of the endosymbionts within each sample, from occupying the majority of the microbial reads (in most samples) to less than 5% of the reads in a few samples (Supplementary Fig. S8). The pronounced turnover of endosymbiont taxa between host populations was mirrored in the Principal coordinate analyses (PCoA): Individuals of the different species from different islands formed separated clusters in PCoA plots and cluster dendrograms for the endosymbiont communities (Fig. 1B, Supplementary Figs. S9 & S10). PCoA plots and cluster dendrograms exhibit a clear pattern of phylosymbiosis, where the divergences between microbiota reflect the phylogenetic distances between the *Ariamnes* hosts. Mantel tests showed a significant correlation between the endosymbiont community beta diversity and the *Ariamnes* (phylo)genetic distances (R>0.6, P<0.001; Table 2 & Supplementary Table S7). Mantel tests similarly showed a significant correlation between the endosymbiont community beta diversity and the geographical distances (R=0.56, p<0.001, Table 2 & Supplementary Table S7), but their correlation coefficients were generally lower that those found with the *Ariamnes* (phylo)genetic distances, suggesting that the different *Ariamnes* host lineages better explain the endosymbiont variation than the geography alone. Within Hawaiʻi Island, endosymbiont communities of *A. waikula* from the same population tend to be more similar than between populations (Supplementary Table S6; *e.g.* individuals from Olaʻa tend to cluster together in Fig. 1B), suggesting a slight population differentiation. However, we found no strong correlation between microbial beta diversity and *A. waikula* (phylo)genetic distances using Mantel tests (Table 2 & Supplementary Table S7).

**Figure 4:**
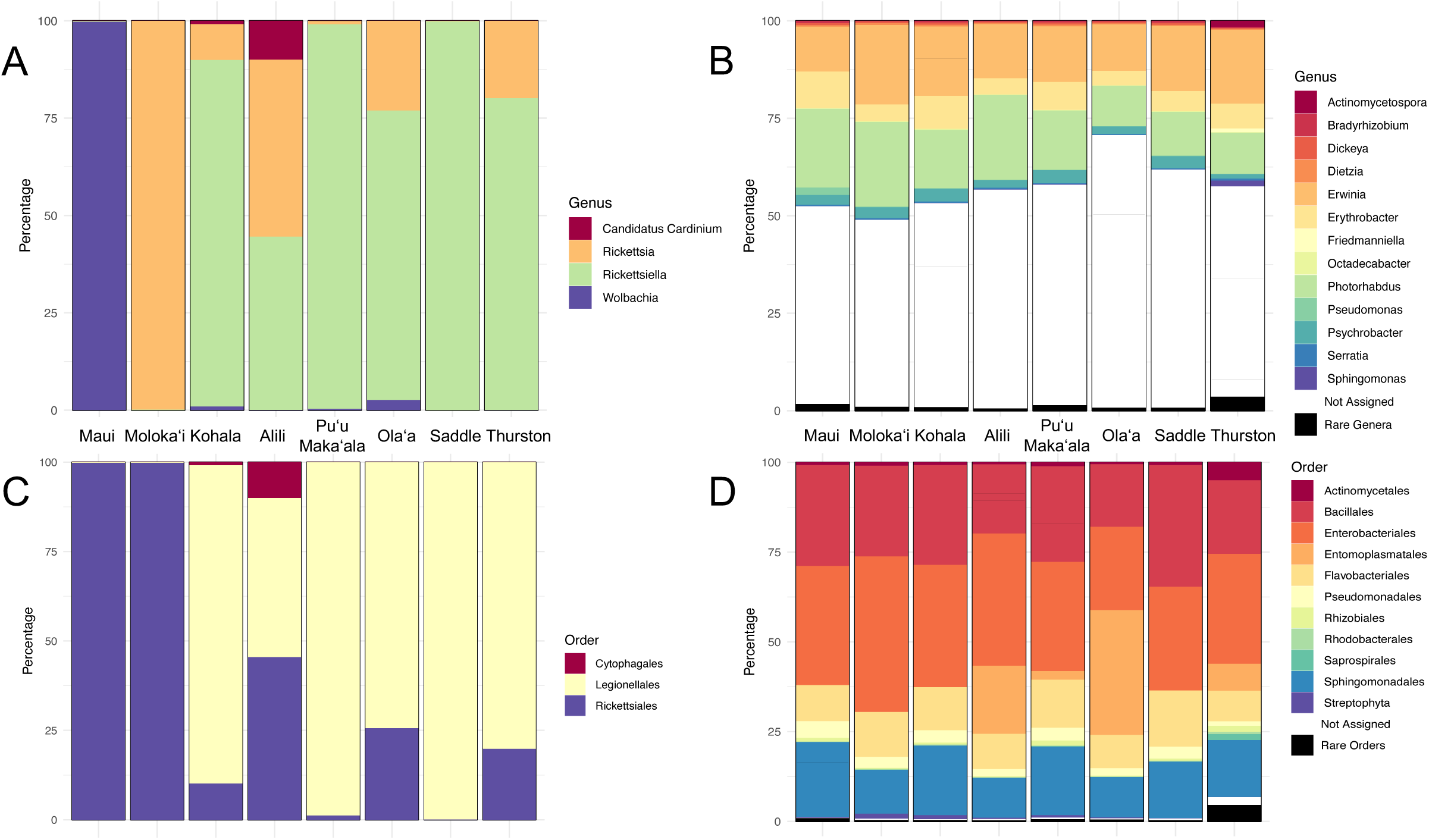
Relative abundances of endosymbionts (A and C) and gut symbionts (B and D) per spider population defined from rarefied Z-OTU table. The panel A-B represents the taxonomic assignations at the genus level, whereas the panel C-D represents the taxonomic assignations at the order level. The relative abundance of each OTU is delimited using the horizontal grey lines and OTUs are colored according to genus. Rare taxonomic assignations (representing less than 1% of the abundance) are merged together. Note that Maui and Molokaʻi correspond to different *Ariamnes* species (*A. melekalikimaka* and *A. n. sp respectively*) whereas the other populations correspond to *A. waikula* on Hawai’i Island. See Table 1 for the number of individual spiders per sampling site.

**Table 2:** Mantel tests comparing the microbial beta diversity dissimilarities to the genetic distances, the phylogenetic distances of their associated *Ariamnes,* and the geographic distances between the sampling sites. Beta diversity dissimilarities were computed on rarefied Z-OTU tables using the Bray-Curtis dissimilarities. Bold values represent significant correlations. Mantel tests were either performed on all populations (between *Ariamnes* species and *A. waikula* populations) or only on *A. waikula* populations.

| Matrix 1 | Matrix 2 | All populations |  | *A. waikula* populations |  |
| --- | --- | --- | --- | --- | --- |
|  |  | Correlation | p-value | Correlation | p-value |
| Whole microbiota beta diversity distances | *Ariamnes* genetic distances | **0.324** | **9.9e-05** | 0.063 | 0.20 |
|  | *Ariamnes* phylogenetic distances | **0.334** | **9.9e-05** | 0.012 | 0.40 |
|  | Geographic distances | **0.303** | **9.9e-05** | 0.053 | 0.20 |
| Endosymbionts beta diversity distances | *Ariamnes* genetic distances | **0.619** | **9.9e-05** | 0.015 | 0.49 |
|  | *Ariamnes* phylogenetic distances | **0.642** | **9.9e-05** | 0.047 | 0.32 |
|  | Geographic distances | **0.567** | **9.9e-05** | 0.108 | 0.169 |
| Gut microbiota beta diversity distances | *Ariamnes* genetic distances | -0.048 | 0.66 | -0.052 | 0.70 |
|  | *Ariamnes* phylogenetic distances | -0.038 | 0.63 | -0.139 | 0.93 |
|  | Geographic distances | 0.007 | 0.42 | -0.084 | 0.78 |

#### Gut microbiota

In contrast to the endosymbionts, the gut microbiota showed a homogeneous taxonomic composition of microbial taxa at the genus level across different *Ariamnes* populations or species (Fig. 4B & Supplementary Fig. S7). This stability of the composition of the gut microbiota was also confirmed at the order level (Fig. 4B & Supplementary Fig. S7) and at the level of the individual spiders (Supplementary Fig. S8). No clear differentiation according to the different species (across islands) or different *A. waikula* populations from Hawai’i Island could be visually detected for the beta diversity of the gut microbiota, either from PCoA decomposition (Supplementary Fig. S9) or hierarchical clustering (dendrogram; Fig. 1C & Supplementary Fig. S10), though there are some slight patterns of clustering by host population. This slight differentiation was confirmed using PERMANOVA: The gut microbiota of individuals from the same population were significantly more similar than the gut microbiota from different populations or species, even within *A. waikula* populations from Hawai’i (R^2^=0.29, P<0.001, Supplementary Table S6). These results indicate that, although the composition of the gut microbiota seems taxonomically similar, there is a slight trend toward microbiota differentiation according to the host populations. However, using Mantel tests we found no significant correlations between the gut microbial community beta diversities and the *Ariamnes* genetic distances, the *Ariamnes* phylogenetic distances, nor the geographical distances (Table 2 & Supplementary Table S7), suggesting that there is likely no pattern of sequential colonization from older to younger volcanoes (stepping stone colonization) in these gut microbial communities in opposition to their spider hosts. We also noticed that besides the slight effect of host populations on the PCoA decomposition (Supplementary Fig. S9) or on the dendrogram of the gut microbiota (Fig. 1C), these gut bacterial communities tend to group into two clusters (Supplementary Fig. S9 and Fig. 1C), with one cluster (“*cluster 1*”) being composed of 14 individual spiders from Alili, Olaʻa, Thurston, and Puʻu Makaʻala. Examination of the details of the differences in gut microbial composition of these two clusters, we found that spider individuals from *cluster 1* were additionally colonized by a single abundant OTU from the order Entomoplasmatales (Supplementary Fig. S11).

Finally, the diversity analyses using phylogenetic metrics showed similar patterns to that obtained with non-phylogenetic metrics (results not shown).

### Simulations

Simulations to assess whether the slow substitutions rates of 16S rRNA can indeed capture signatures of change among host-associated microbes in recently evolving lineages (<2MY) showed that we can recover a significant phylosymbiotic pattern, i.e. a significant correlation between the microbial beta diversity and the *Ariamnes* phylogenetic distances, even when only 10% of the bacterial OTUs were transmitted and 90% of the remaining OTUs were evolving independently from the *Ariamnes* phylogeny (Figure S12). When >=75% of the bacterial OTUs were transmitted, we always recovered a strong and significant phylosymbiosis (Figure S12). This suggested that, despite the slow evolution of the 16S rRNA gene, we should be able to detect microbiota differentiation if some of the *Ariamnes*-associated bacteria diverged separately in the different populations, whether because of vertical transmission or because of vicariance events in the environmental pools of available bacteria. Moreover, simulations of the bottlenecks in microbial diversity during island colonizations showed that we can recover a signal in the microbial alpha diversity (a decrease of diversity in the youngest sites) even if the bottleneck has impacted only a small proportion of the bacterial OTUs (<10%; Figure S13).

## Discussion

In this study, we examined the early stages of diversification of Hawaiian *Ariamnes* spiders in concert with the different components of their associated microbial community on the youngest island of the Hawaiian archipelago. Our results show a strong effect of isolation by distance in the host and contrasting patterns of community composition among both the endosymbionts and the gut microbiota they harbor, suggesting eco-evolutionary distinctiveness of the host and each of the two microbial components.

### Isolation by distance explains Ariamnes diversification

Our analyses suggest that the nascent diversification of *A. waikula* is mainly the result of isolation by distance. The different populations of *A. waikula* show a predictable pattern of colonization across Hawaiʻi Island from the oldest (Kohala) to the youngest (MLKS) sites, with little genetic structure between the geographically proximal MLKS sites of Olaʻa, Puʻu Makaʻala, and Thurston (PCA, Supplementary Fig. S5; FST, Supplementary Table S3; Admixture, Fig. 2B). Our analyses suggest either that gene flow is still occurring between these MLKS populations, or that the recency of colonization means that there has been insufficient time for differentiation in this area (Roderick et al. 2012). Interestingly, we did not find a significant increase of genetic diversity with volcano age, as expected with successive founding events. These patterns may become more apparent with additional loci or more extensive sampling.

Patterns of genetic differentiation suggest a potentially complicated course of colonization across Hawaiʻi Island for *A. waikula*. Although Kohala, which represents the oldest volcano, was consistently characterized as distinct from other populations in PCA, phylogenetic, and admixture analyses, both Saddle and Alili were also largely distinct. These patterns are potentially explained by a shifting mosaic population structure (Carson et al 1990) with multiple colonization events, historical connectivity, or ongoing admixture. However, sampling of additional sites, individuals, and species will be required to disentangle the historical sequence of events explaining the relationship between these sites. Our results are consistent with other population genetic studies on Hawaiʻi Island showing strong differentiation and potential speciation of lineages on the different volcanoes within the island (Funk & Wagner 1995; Holland & Cowie 2007; Muir & Price 2008; Croucher et al. 2012; Goodman et al. 2019; Eldon et al. 2019; Blankers et al. 2018). Interestingly, spiders on the mainland would not show this isolation by volcano pattern, because they do efficiently disperse by ballooning; here, our data, in contrast, suggest that the species does not efficiently balloon anymore in this island system (Gillespie et al 2012).

### Major endosymbiont changes after potential colonization event

We found major shifts among the dominant endosymbiont genera across the different islands and *Ariamnes* species: Species of *Ariamnes* from the two geologically oldest sites included in the study (Molokaʻi and Maui) each harbor a unique endosymbiont genus (*Rickettsia* and *Wolbachia* respectively) while the different populations of *A. waikula* on the youngest Hawaii Island all have a mix of endosymbionts dominated by the genus *Rickettsiella*. Most spider individuals were colonized by a single endosymbiont OTU as previously observed in *Argiope* spiders (Sheffer et al. 2020), but there were also a few individuals that were co-colonized by both *Rickettsiella* and *Rickettsia*. Although such endosymbiont coexistence has been previously observed (*e.g.* Mouton et al. (2004) found up to 3 coexisting *Wolbachia* strains in some wasps), we cannot exclude the possibility that one of these endosymbionts may have come from a prey item that had just been eaten before sampling (Kennedy et al. 2020). The overall shifts in endosymbionts across *Ariamnes* species likely contribute to the congruence between the host phylogeny and the dendrogram of microbiota dissimilarity - a pattern referred as phylosymbiosis (Brooks et al. 2016; Lim & Bordenstein, 2020). The strong conservatism of the dominant endosymbiont within each island is consistent with the maternal inheritance and vertical transmission of the endosymbionts (Duron et al. 2008) and the fact that it generates phylosymbiosis has been similarly observed in other arthropods (e.g. a *Camponotus* ant system and its associated endosymbiont *Candidatus Blochmania* (Degnan et al. 2004)). Therefore, the endosymbiont component of the microbiota reflects the overall evolutionary relationships of the *Ariamnes* host. However, from our study, we cannot say whether endosymbionts played a role in promoting the genetic isolation of *A. waikula* populations; this hypothesis would require experimental manipulation of microbial communities within different host populations.

The observed turnover of dominant endosymbiont genera between islands and host populations suggests that endosymbiont horizontal transmission happens quickly. Previous studies have also shown frequent horizontal transfers in the *Wolbachia* strains infecting arthropods, this pattern being more common than co-speciation events (Baldo et al., 2008; Stahlhut et al. 2010). In particular, Stahlhut et al. (2010) showed that hosts sharing the same ecology were likely to horizontally exchange *Wolbachia* strains. In contrast, a more recent study reported lower turnover rates over time (7 Myr on average) for the bacterial genus *Wolbachia* colonizing arthropods in French Polynesia and found that *Wolbachia* colonization did not seem related to recent events of isolation mediated by island structure (Bailly-Bechet et al. 2017). Given that the islands used in the current study, and thus the spider populations studied here, are separated by <2 Myr (Gillespie et al. 2018), the more rapid turnover in *Ariamnes* endosymbiotic communities (compared to *Wolbachia* in the French Polynesian arthropods) appears to be linked to the geological formation of islands. Therefore, it seems to exist a range of endosymbiont dynamics, from conserved and stable host associations (Degnan et al. 2004) to labile and horizontally acquired (Baldo et al., 2008; Stahlhut et al. 2010; this study). For *Ariamnes* spiders, including *A. waikula* (Kennedy et al. 2018), predation on other arthropods and cannibalism are particularly important (Whitehouse et al. 2002), providing a potential avenue for endosymbiont horizontal transfer (Su et al. 2019) and may explain the high endosymbiont turnover observed in this system. However, from our current analyses, because we did not study the phylogenetic history of individual microbial lineages, we cannot date the events of endosymbiont acquisition. In particular, we cannot separate a scenario of recent horizontal shifts of the endosymbionts, where endosymbionts were very recently acquired from the environment, from a scenario of older shifts, where endosymbionts were acquired during colonization events and were thus synchronous with speciation events or subject to priority effects. Testing this will require modeling host-microbiota evolution, which is currently challenging on such short evolutionary timescales (Groussin et al. 2017; Perez-Lamarque & Morlon 2019).

Several previous studies have suggested that endosymbionts such as *Rickettsia* reduce the behavioral propensity for ballooning dispersal in spiders (Goodacre et al. 2009). Hawaiian *Ariamnes* have never been shown to balloon and exhibit high local endemism, a common phenomenon in organisms endemic to remote islands, generally attributed to distinct local selective pressures (Gillespie et al 2012). The possibility that such a reduction in dispersal may be associated with bacterial infection is intriguing. Experimental studies will be needed to test the hypothesis.

### Similarity of the gut microbiota

Across all spider populations, we observed a conserved gut microbiota: Over approximately one million years of divergence, the composition seems to have remained stable and does not reflect the *Ariamnes* genetic structure. The core lineages at the generic level are the same as those typically found in the gut microbiota of other arthropods (Engel & Moran 2013), including spiders (Kennedy et al. 2020, Sheffer et al. 2020). In addition, although we detected slight but significant differences in the gut microbiota between the different spider species and between the *A. waikula* populations within Hawai’i Island, these patterns of differentiation were not associated with *Ariamnes* (phylo)genetic structure, volcano age, nor the geographical distances between the sampling sites. Most of the variation in the gut microbiota observed using PCoA or cluster dendrograms (Fig. 3C and Supplementary Fig. S9) was actually explained by a single OTU from the order Entomoplasmatales that was present in high abundance in many *A. waikula* individuals across different populations (Supplementary Fig. S8). Given that Entomoplasmatales are arthropod pathogens (Gasparich, 2014), the observed variation across *Ariamnes* gut microbiota may simply reflect the local acquisition from the environment, decorrelated from the host (phylo)genetic structure.

We found that both alpha and beta diversity of gut microbes was similar across the different islands and populations, decoupled from the spider’s phylogeny. This absence of a correlation with the host’s phylogenetic structure suggests that the gut microbiota is likely acquired from the spider’s environment, most probably through the spider’s diet (Kennedy et al. 2020; Zhang et al. 2018). The pattern of slight differentiation according to the host populations could either be due to gradual changes in the host-filtering process (Mazel et al. 2018; see Introduction), or only reflect the local variations of the bacterial pool available in the host environment.

One could argue that the absence of strong differentiation in the microbial communities could come from the slow evolution of the 16S rRNA marker which may not provide the needed resolution to show differences at short timescales (<2 Myr). However, given that we used Z-OTU clustering (0% radius OTUs) that can detect any variation in the 16S sequences and that the 16S gene of symbiotic bacteria is estimated to diverge at least 1-2% per 50 Myr on average (Ochman et al., 1999), our simulations demonstrated that we should expect to have some variation in a portion of the 16S sequences of the transmitted bacteria. Thus, we should expect to detect microbial differentiation (by comparing their beta diversity) and a resulting pattern of phylosymbiosis (Sanders et al. 2014, Perez-Lamarque & Morlon, 2019). In particular, our simulations demonstrated that even 10% transmission of the microbial OTUs can be sufficient to generate a significant pattern of phylosymbiosis within such a short amount of time. In addition, if the *Ariamnes*-associated microbial communities experienced bottlenecks at colonization events, we expect filtering of the preexisting (genetic or strain) diversity that has accumulated for more than 2 Myr, and this should not be affected by the slow evolution of the 16S rRNA gene. Here again, this expectation was supported by the simulations which showed that diversity bottlenecks in the microbial communities resulting from a stepping stone colonization, could readily be detected and was not sensitive to the short timescales of the *A. waikula* divergences.

The overall similarity of the gut microbiota at the generic level has been shown in other systems, such as recently diversified *Anolis* lizards (Ren et al. 2016) and *Cephalotes* turtle ants (Sanders et al. 2014). Such similarity likely conserves the functional properties of the gut microbiota (Muegge et al. 2011), which could be particularly advantageous for arthropods which often rely on gut symbionts to complement imbalanced nutrition (Engel & Moran 2013) or detoxify toxin-rich diets (Adams et al. 2013). It is unclear whether *Ariamnes* spiders, and predatory arthropods in general, also rely on their microbiota for various functions. A recent study suggested that, the spider microbiota can be mainly derived from the prey items, without any apparent functions (Kennedy et al., 2020). If the same applies for the Hawaiian *Ariamnes*, the conserved gut microbiota of *A. waikula* may reflect its conserved niche, with the diet presumably similar despite changes in geographical range.

## Conclusion

Organisms host unique and large microbial diversities, and it is likely that the different components of the microbial community experience different ecological and evolutionary dynamics. In this study, we demonstrated a dichotomy between the endosymbiont communities and the gut microbiota: For the host Hawaiian *Ariamnes* spiders, abiotic factors, in particular isolation by distance along the volcano chronosequence, appear to have the strongest influence in explaining divergence patterns. The evolutionary history of the hosts also strongly impacts the colonization of endosymbionts, which show clear shifts in their composition between species on different islands. However, we find little differentiation in endosymbionts across populations of *A. waikula* within Hawaʻi Island, despite the strong genetic structure of the host. In contrast to the differences we find between taxa in the endosymbiont composition, the composition of the gut microbiota is overall similar across both species and populations despite strong geographic isolation of their hosts. Our results stress the high heterogeneity within the different components of the *Ariamnes* holobiont that likely did not act as a single homogenous unit of selection. Investigating the composition of microbiota through a broader sampling across different ecomorphs and islands would significantly contribute to our understanding of the dynamics of host-microbe interactions as it pertains to rapid speciation events.

## Acknowledgements

We would like to acknowledge support and assistance from the following: The permit processing and access to different reserves and private land was possible thanks to Steve Bergfeld (DOFAW Big Island), Pat Bily (TNC Maui), Tabetha Block (HETF), Pomaikaʻi Kaniaupio-Crozier (Maui Land and Pineapple), Lance DaSilva (DOFAW Maui), Charmian Dang (NAR), Melissa Dean (HETF), Betsy Gagne (NAR), Lisa Hadway (DOFAW Big Island), Cynthia King (DLNR), Russell Kallstrom (TNC Molokaʻi), Joey Mello (DOFAW Big Island), Ed Misaki (TNC Molokaʻi). For support and advice in the lab, we are very grateful to Lydia Smith (Evolutionary Genetics Lab, Museum of Vertebrate Zoology, UC Berkeley), Shana McDevitt (Vincent J. Coates Genomics Sequencing Laboratory, QB3, UC Berkeley), and Anna Sellas (California Academy of Sciences, San Francisco). Samples were provided by Susan Kennedy and Andrew Rominger. We thank Benjamin Peter, Thorfinn S. Korneliussen, and Line Skotte for advice on the analyses. L.E.B was supported by the Netherlands Organisation for Scientific Research VENI #863.14.020. B.P.L was supported by a master fellowship from the École Normale Supérieure of Paris. H.K. was supported by a postdoctoral fellowship by the German Research Foundation (DFG). Support for the project was provided by the NSF DEB 1241253 to R.G.G.

## Data Accessibility

The pipelines used for processing ddRAD seq data are available in https://github.com/CGRL-QB3-UCBerkeley/RAD. All scripts used to analyze the Ariamnes microbiota are available in https://github.com/BPerezLamarque/Scripts. The raw data can be found on dryad (https://datadryad.org) DOI: https://doi.org/10.5061/dryad.nzs7h44qj

## Author Contributions

R.G.G and H.K. conceived of the study. E.A. and B.P-L. performed DNA extractions. C.C. and L.E.B. performed ddRAD lab work. E.A., K.B., and T.L. performed ddRAD genetic analyses. B.P-L. and H.K. performed lab work and analyses for the microbial component of the study. E.A., B.P-L., H.K., and R.G.G wrote the manuscript. All authors edited and approved the manuscript before final submission.

## Supplementary Figures

**Figure S1:**
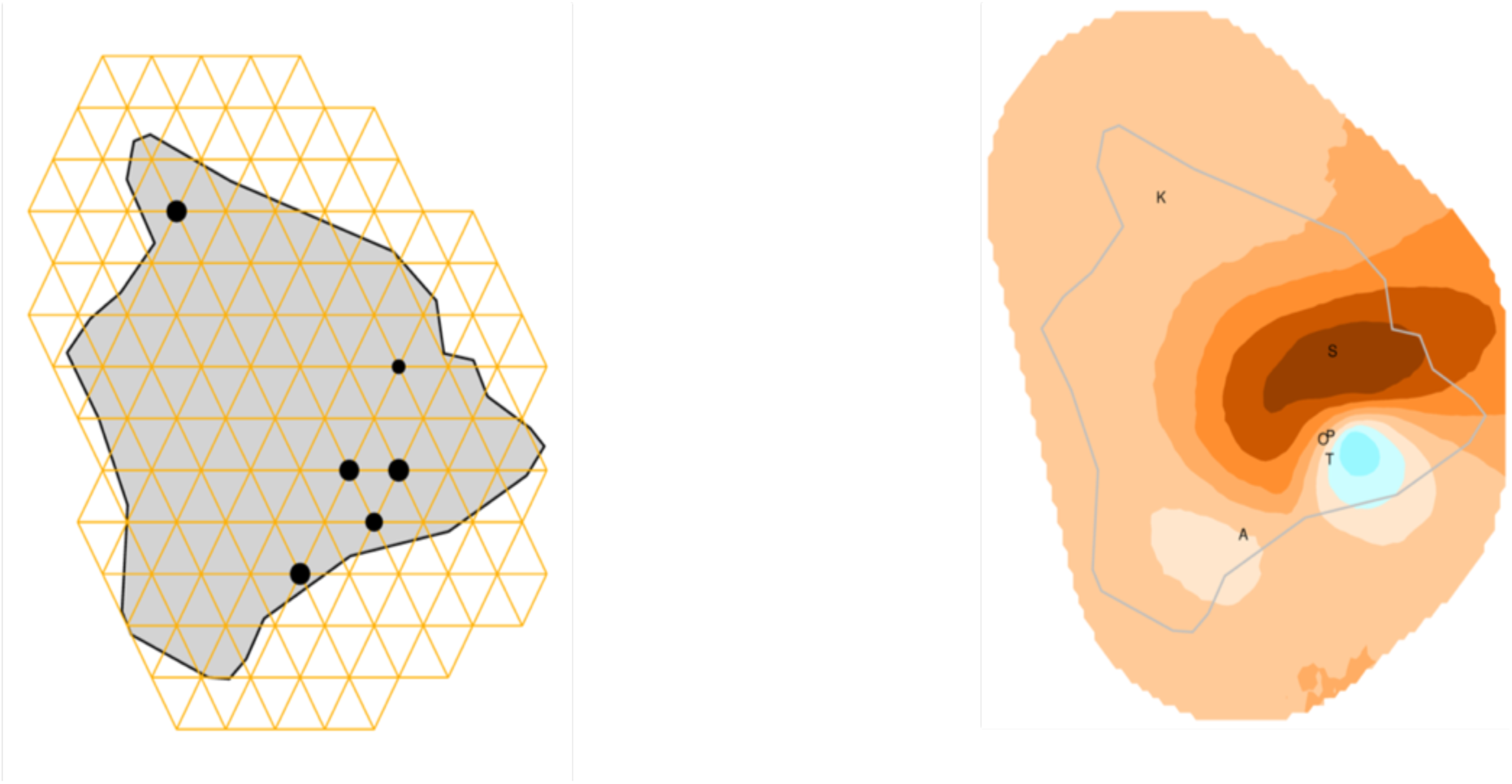
Grid system used for the EEMS analyses on Hawaiʻi Island (left) and corresponding results (right) A: Alili, K: Kohala, O: Olaʻa , P: Puʻu Makaʻala, S: Saddle, T: Thurston

**Figure S2.**
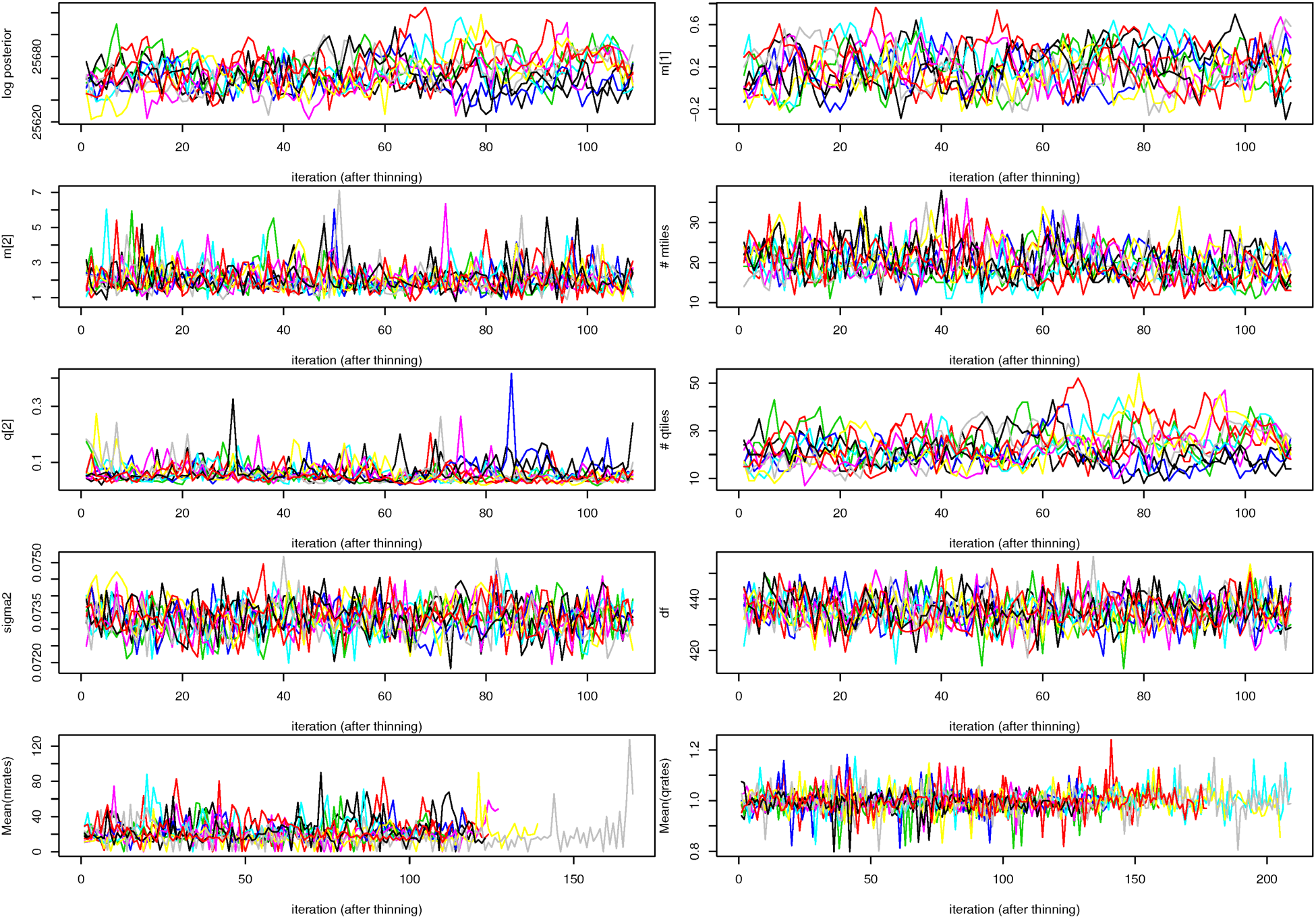
Traces of the MCMC runs of the 10 replicates to assess convergence. Only individuals of the gold ecomorph from Hawaiʻi Island were used here.

**Figure S3:**
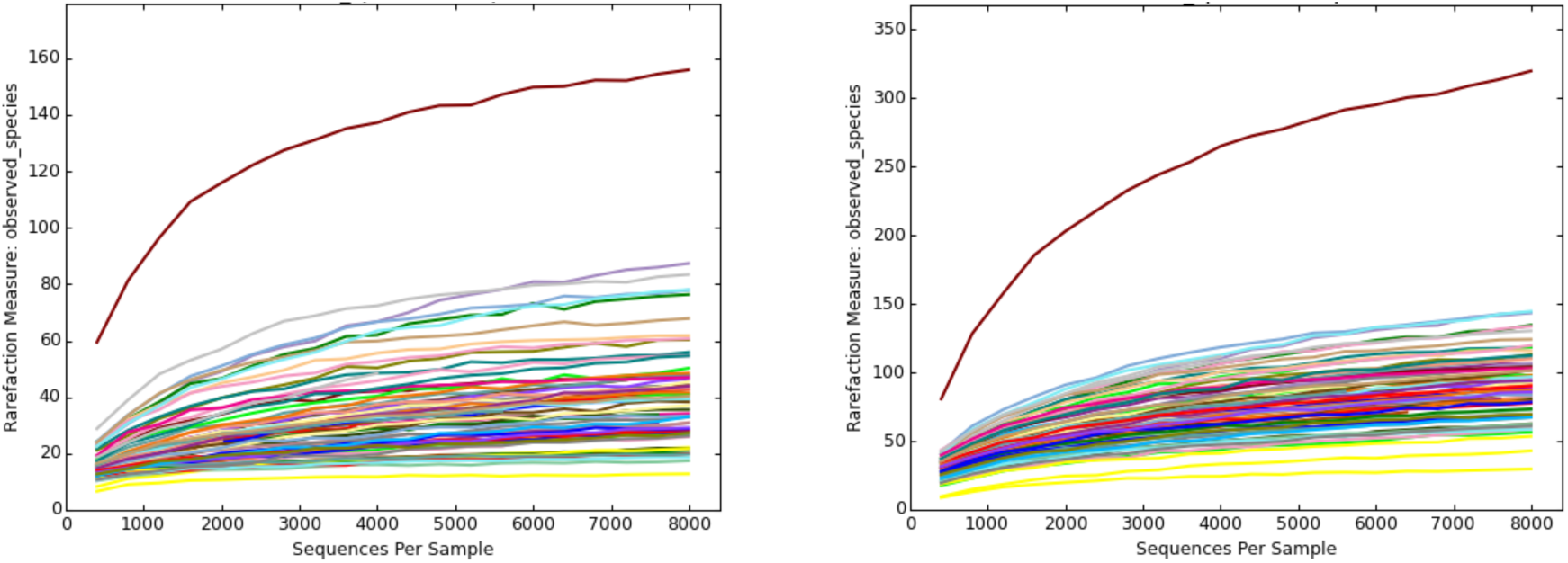
Rarefaction curve of the whole microbiota: number of observed OTU in each sample according to the number of rarefied reads per sample. Results of OTU at 97% are presented on the left, whereas Z-OTU are on the right. Rarefactions are performed 20 times independently and the mean value is plotted here. Note that one sample (from Thurston) present an unexpectedly high diversity compared to other samples.

**Figure S4:**
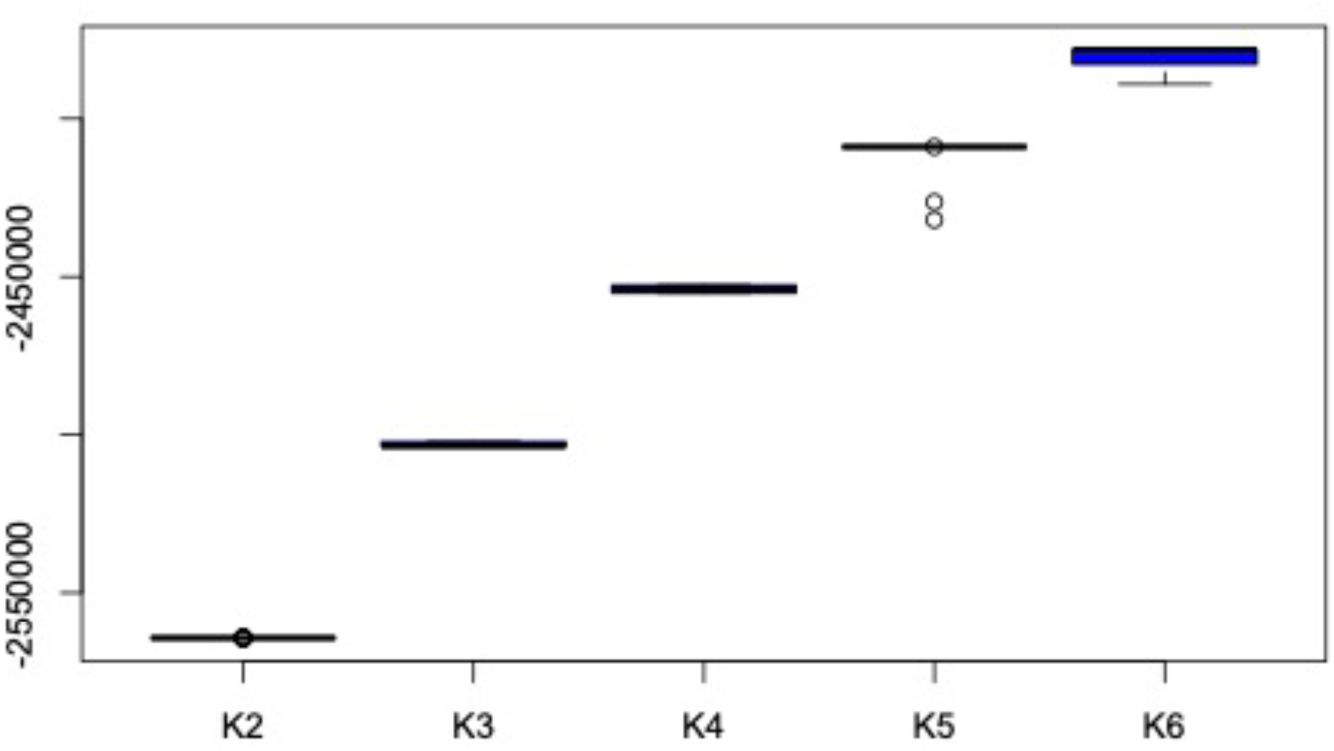
Log likelihood for 50 iterations of NGSAdmix K values 2-6 for *A. waikula* from Hawai’i Island.

**Figure S5:**
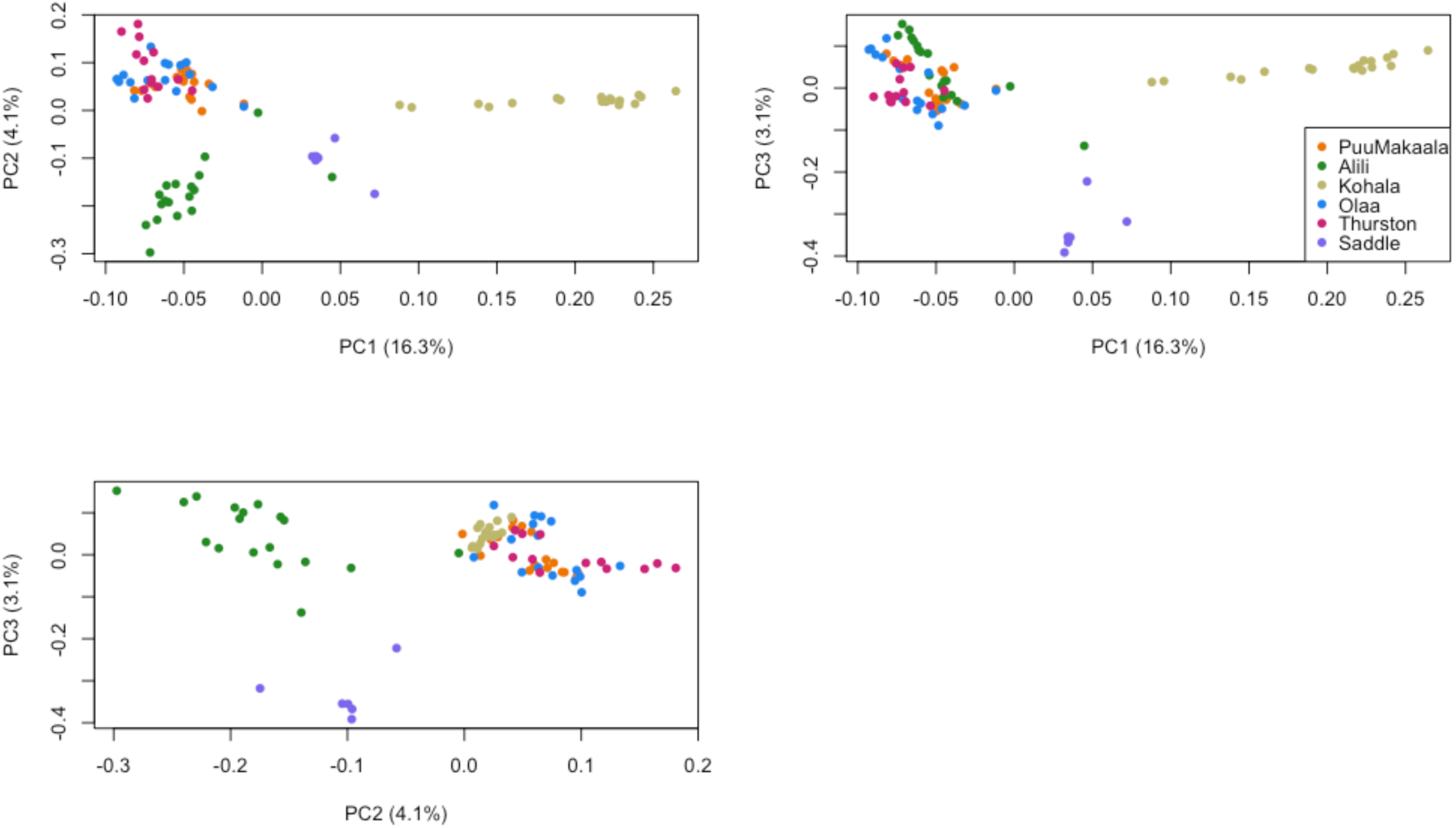
PCA analyses of ddRAD data of *A. waikula* from Hawai’i Island

**Figure S6:**
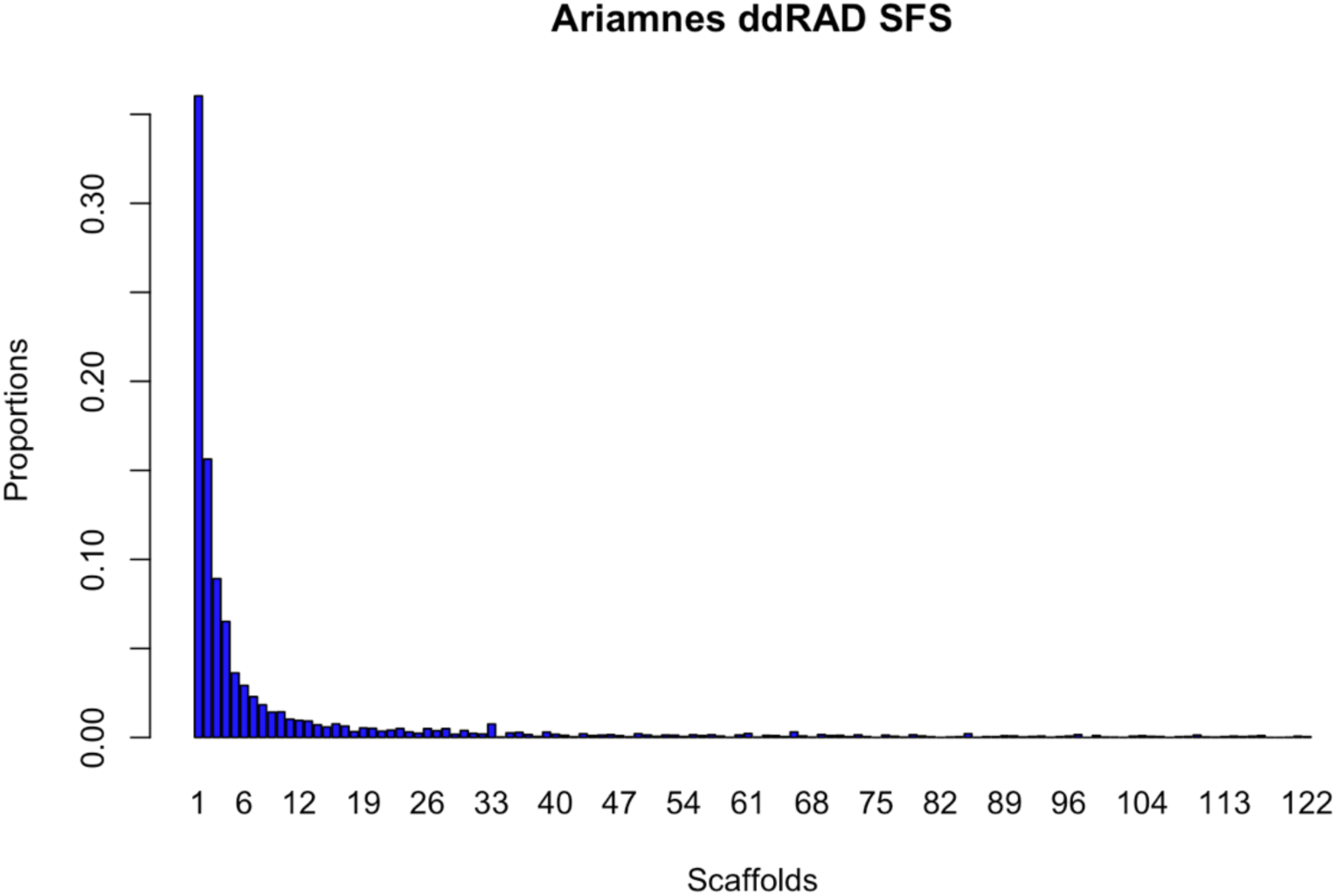
Site frequency spectrum of all *Ariamnes* ddRAD data after filtering.

**Figure S7:**
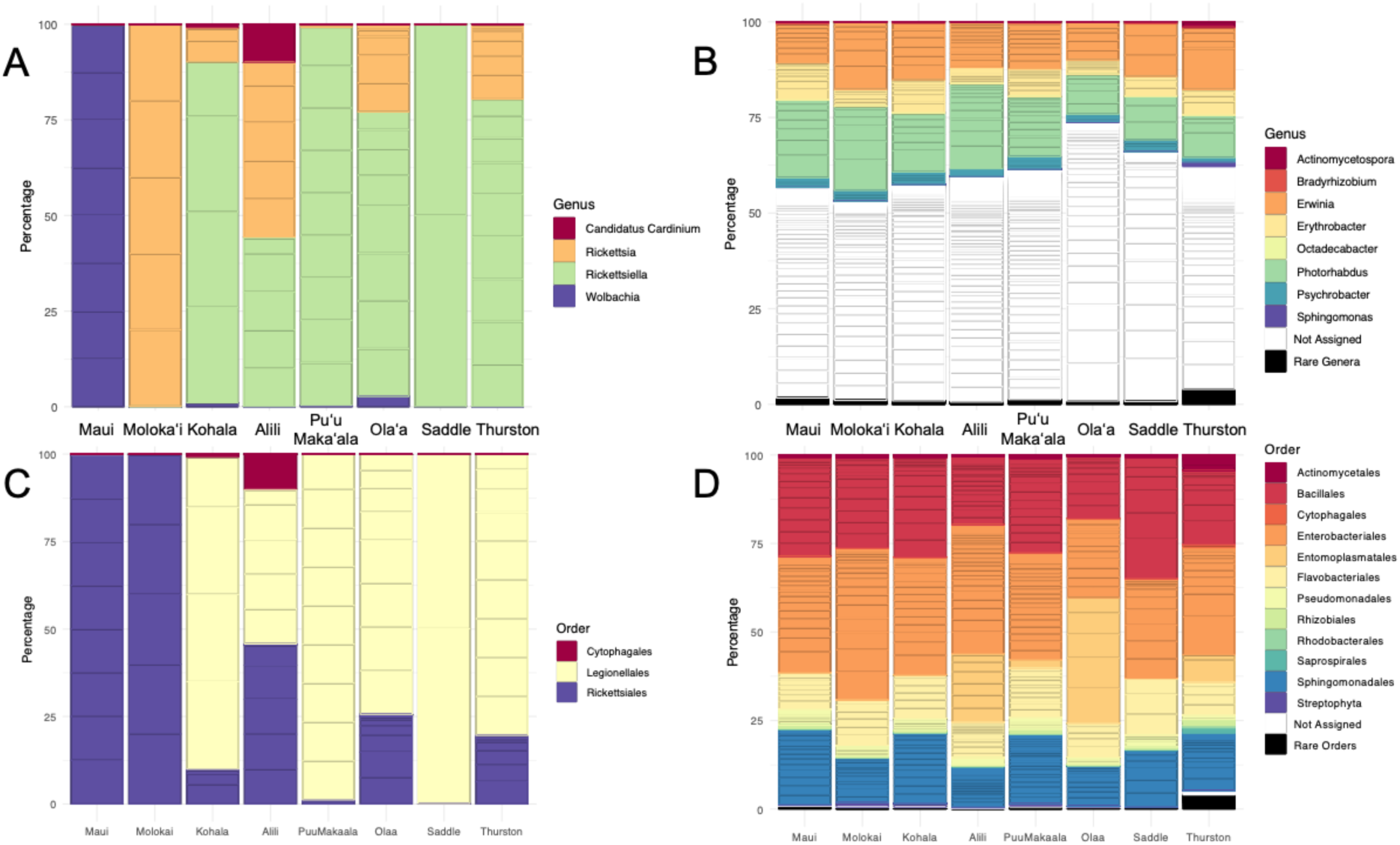
Relative abundances of endosymbionts per population (A and C) and gut symbionts per spider population (B and D) defined from rarefied 97% OTU table. The panel A-B represents the taxonomic assignations at the genus level, whereas the panel C-D represents the taxonomic assignations at the order level. The relative abundance of each OTU is delimited using the horizontal grey lines and OTUs are colored according to genus/order. Rare taxonomic assignations (representing less than 1% of the abundance) are merged together. Note that Maui and Molokaʻi correspond to different *Ariamnes* species (*A. melekalikimaka* and *A. n. sp respectively*) whereas the other populations correspond to *A. waikula* on Hawai’i Island. See Table 1 for the number of individual spiders per sampling site.

**Figure S8:**
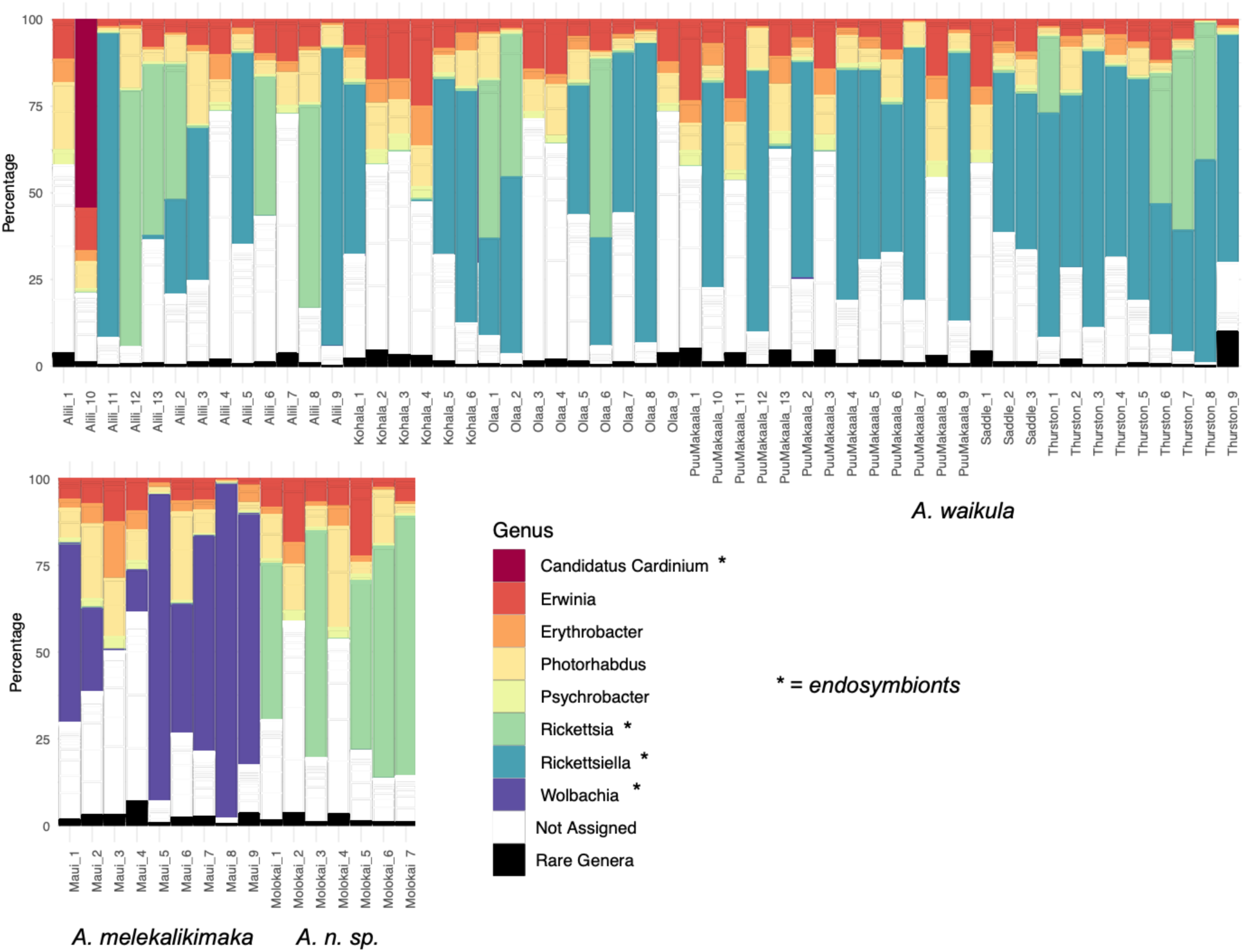
Relative abundances of the microbial community (both endosymbionts and gut symbionts) per spider individual from rarefied Z-OTU table. The relative abundance of each OTU is delimited using the horizontal grey lines and OTUs are colored according to genus. Rare taxonomic assignations (representing less than 0.5% of the abundance) are merged together. Note that Maui and Molokaʻi correspond to different *Ariamnes* species (*A. melekalikimaka* and *A. n. sp respectively*) whereas the other populations correspond to *A. waikula* on Hawai’i Island.

**Figure S9:**
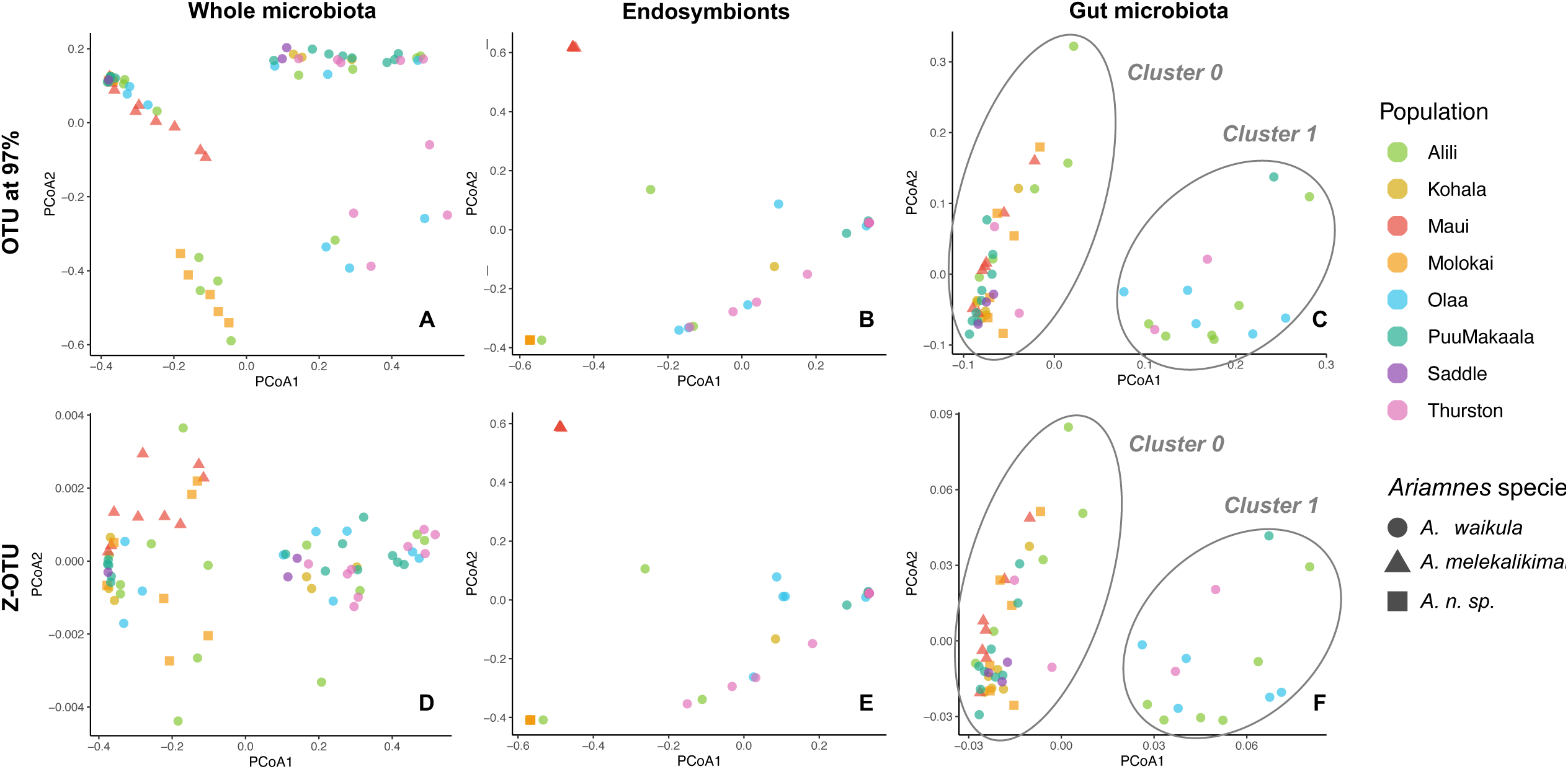
Principal coordinate analyses (PCoA) decomposition of the microbial communities according to the type of OTU clustering (OTU at 97% or Z-OTU) and the type of microbial communities (whole microbiota, endosymbionts, or gut microbiota). Note that Maui and Molokaʻi correspond to different *Ariamnes* species (*A. melekalikimaka* and *A. n. sp respectively*) whereas the other populations correspond to *A. waikula* on Hawai’i Island. We noticed the presence of two clusters in the PCoA of the gut microbiota that are not explained by the identity of the host species or population: these two clusters (cluster 0 and cluster 1) are highlighted in grey.

**Figure S10:**
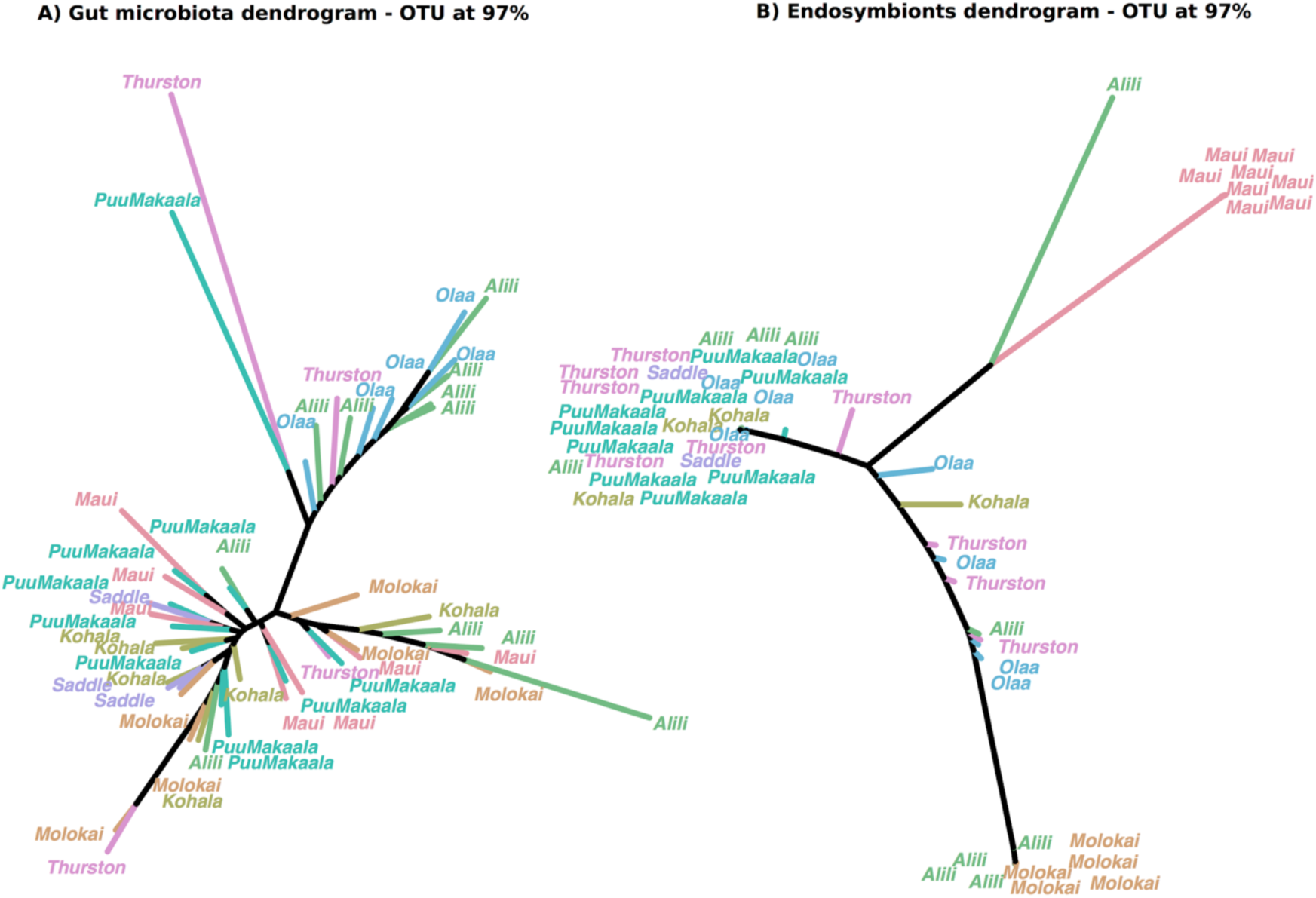
(A) Microbiota dendrograms reconstructed from the gut microbiota and (B) the endosymbiont community, for the OTU at 97%. Note that Maui and Molokaʻi correspond to different *Ariamnes* species (*A. melekalikimaka* and *A. n. sp respectively*) whereas the other populations correspond to *A. waikula* on Hawai’i Island.

**Figure S11:**
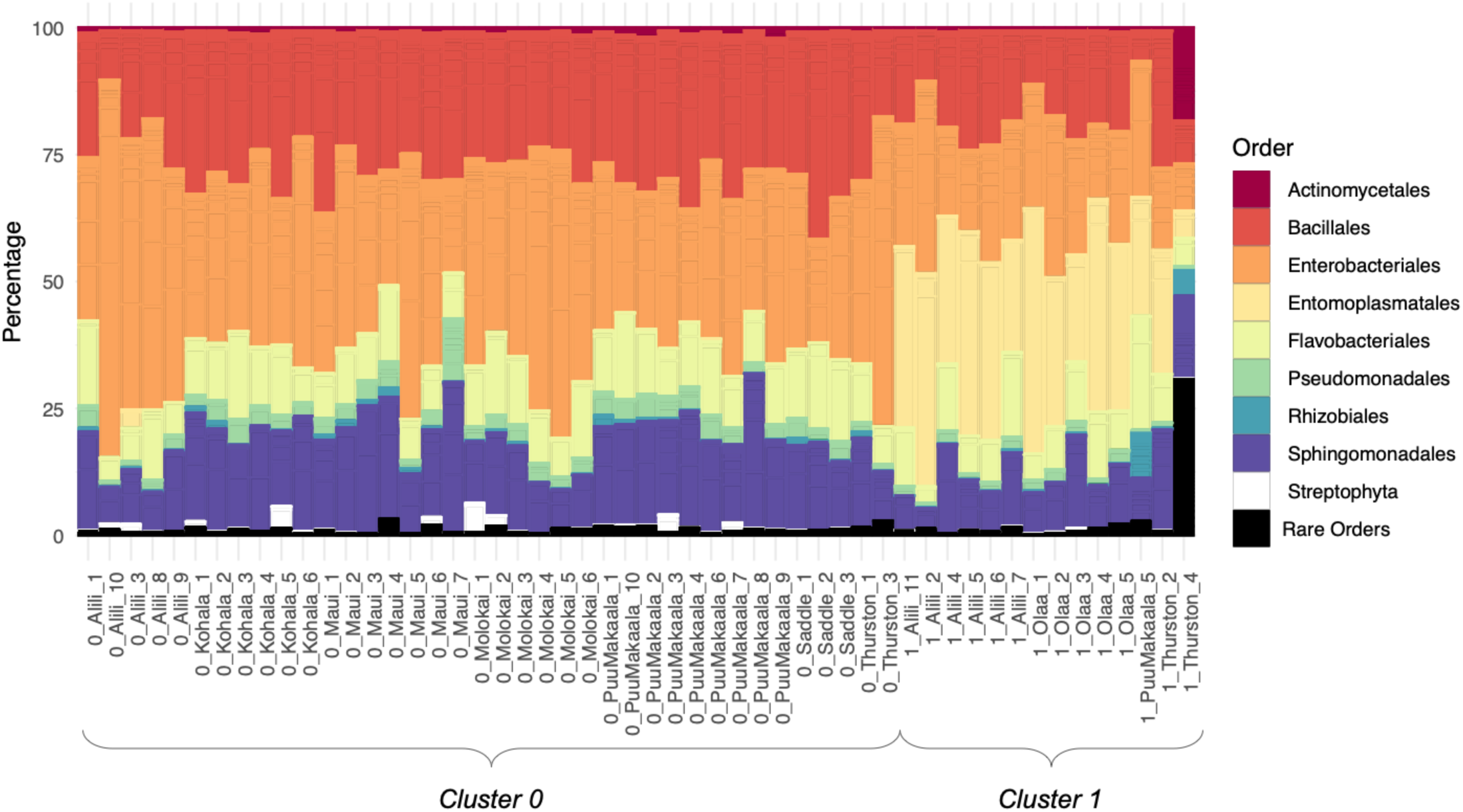
Relative abundances of the gut microbial community per spider individual from the rarefied Z-OTU table. The relative abundance of each OTU is delimited using the horizontal grey lines and OTUs are colored according to order. Rare taxonomic assignations (representing less than 0.5% of the abundance) are merged together. Note that Maui and Molokaʻi correspond to different *Ariamnes* species (*A. melekalikimaka* and *A. n. sp respectively*) whereas the other populations correspond to *A. waikula* on Hawai’i Island. Samples were sorted by cluster (cluster 0 and cluster 1), which emerged in the PCoA decomposition of the gut microbiota (see Fig. S9).

**Figure S12:**
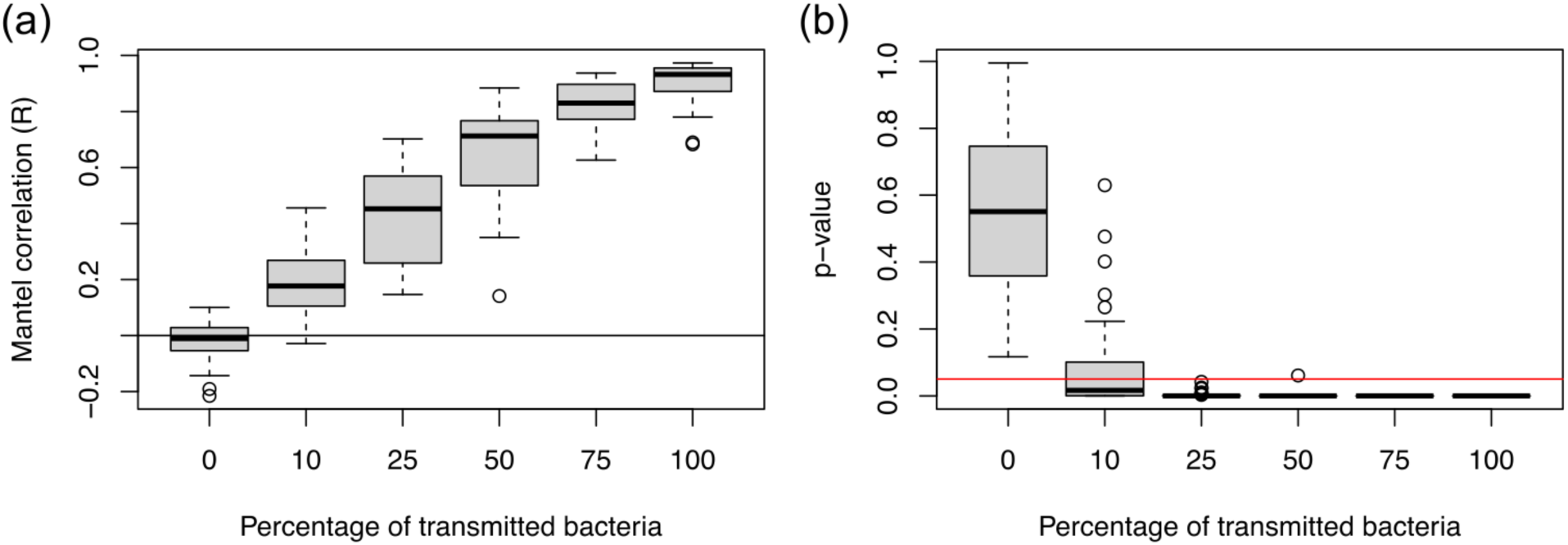
Pattern of phylosymbiosis as a function of the simulated percentage of transmitted bacteria: For each percentage of vertically transmitted bacteria (0, 10, 25, 50, 75, or 100%), 30 mock microbiota communities, each of 100 OTUs, associated with *Ariamnes* spiders were simulated. Mantel tests between the *Ariamnes* phylogenetic distances and the simulated microbial beta diversity (Bray-Curtis) show (a) the Pearson correlation values (R) of the Mantel tests and (b) the corresponding p-values (obtained with 10,000 permutations – the red horizontal line indicates the significance threshold of 0.05, i.e. below this line there is a significant pattern of phylosymbiosis). Note that the results are qualitatively similar when using UniFrac distances instead of Bray-Curtis dissimilarities.

**Figure S13:**
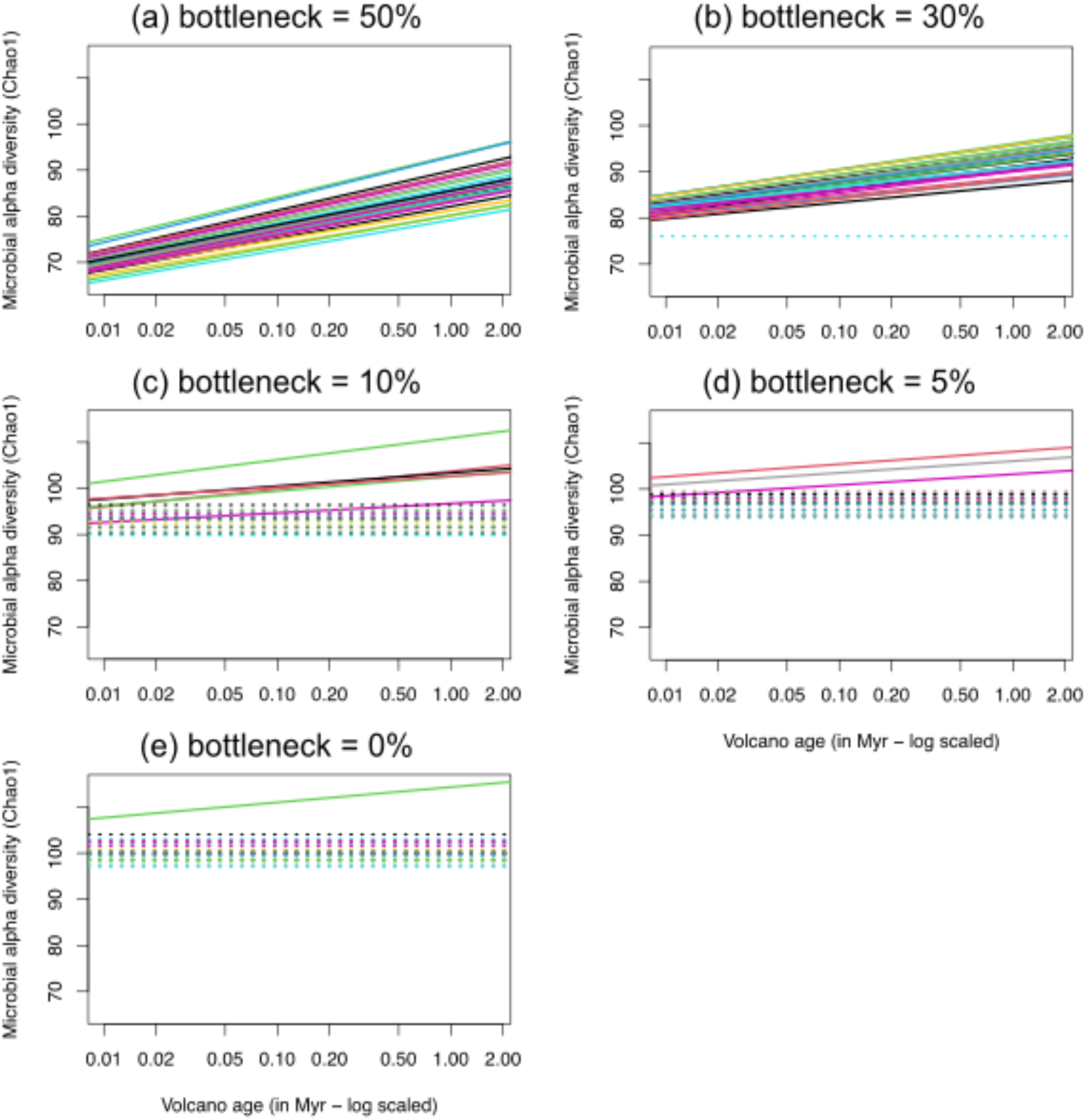
Presence of a bottleneck in *Ariamnes* microbial alpha diversities when simulating a bottleneck in the environmental pools of available microbes. We simulated a diversity bottleneck along the island chronosequence (between the older Molokaʻi-Maui islands and the recent Hawaiʻi Island) and randomly assembled mock *Ariamnes* microbiota from the pool of available microbes (see Methods). For each percentage of microbial OTUs experiencing the bottleneck (0, 5, 10, 30, or 50%), we replicated 30 mock microbiota communities and tested whether the alpha diversities (Chao1 index) of the *Ariamnes* microbiota increase with the volcano age (along the chronosequence) using linear mixed models. Plots show solid lines when the relationship was significant (p-value<0.05), dotted lines when not significant.

## Supplementary Tables

**Table S1:** Specimen collection detail for population genetic and microbial samples.

| Species | Eco-morph | Island | Population | Volcano (age) | Substrate age (years) | Latitude | Longitude | Collector | Date | Individuals (ddRAD) | Microbiota |
| --- | --- | --- | --- | --- | --- | --- | --- | --- | --- | --- | --- |
| *A. waikula* | Gold | Hawai'i | Kohala | Kohala, 0.43 | 300,000 | 20.1048 | -155.7242 | Rominger | 2014-01-12 | 18 | 6 |
| *A. waikula* | Gold | Hawai'i | Saddle | Mauna Loa, Kea 0.01-0.38 | 4,000 | 19.6746 | -155.3316 | Rominger | 2014-01-07 | 6 | 3 |
| *A. waikula* | Gold | Hawai'i | Puu Makaala | Kilauea, 0.004 | 11,000 | 19.3748 | -155.2202 | Rominger | 2014-01-05 | 18 | 13 |
| *A. waikula* | Gold | Hawai'i | Thurston | Kilauea, 0.004 | 600 | 19.4078 | -155.2388 | Rominger | 2014-01-08 | 13 | 9 |
| *A. waikula* | Gold | Hawai'i | Ola'a | Kilauea, 0.004 | 7,500 | 19.4530 | -155.2466 | Rominger | 2014-01-09 | 16 | 11 |
| *A. waikula* | Gold | Hawai'i | Alili | Mauna Loa, 0.01 | 20,000 | 19.2309 | -155.5134 | Rominger | 2014-01-04 | 18 | 13 |
| *A. hiwa* | Brown | Hawai'i | Puu Makaala | Kilauea, 0.004 | 11,000 | 19.3748 | -155.2202 | Rominger | 2014-01-05 | 1 | N/A |
| *A. hiwa* | Brown | Hawai'i | Thurston | Kilauea, 0.004 | 600 | 19.4078 | -155.2388 | Rominger | 2014-01-08 | 1 | N/A |
| *A. meleka-likimaka* | Gold | W. Maui | Puu Kukui | Puu Kukui, 1.3 | 1,500,000 | 20.9194 | -156.5972 | Gillespie | 2012-06-29 | 16 | 9 |
| *A. n. sp* | Gold | Moloka'i | Kamakou | Moloka'i, 1.8 | 1,400,000 | 21.1108 | -156.8906 | Gillespie | 2012-06-23 | 16 | 7 |

**Table S2:** Specimen collection detail for population genetic and microbial samples.

| Island | Population | Name ddRAD | Name microbiota | SeqID | Pool | Sample ID |
| --- | --- | --- | --- | --- | --- | --- |
| Hawai'i | Pu'u Maka'ala | 1 | 83 | RGCC001A_S25_AACCA | A2 | AJR923 |
| Hawai'i | Pu'u Maka'ala | 2 | NA | RGCC001A_S25_AAGGA | A8 | AJR913 |
| Hawai'i | Pu'u Maka'ala | 3 | 82 | RGCC001A_S25_ACACA | A10 | AJR948 |
| Hawai'i | Pu'u Maka'ala | 4 | NA | RGCC001A_S25_ACTGG | A13 | AJR956 |
| Hawai'i | Pu'u Maka'ala | 5 | 70 | RGCC001A_S25_ACGGT | A12 | AJR955 |
| Hawai'i | Pu'u Maka'ala | 6 | 65 | RGCC001A_S25_ACTTC | A14 | AJR964 |
| Hawai'i | Pu'u Maka'ala | 7 | 47 | RGCC001A_S25_AGCTA | A9 | AJR915 |
| Hawai'i | Pu'u Maka'ala | 8 | NA | RGCC001A_S25_ATACG | A15 | AJR965 |
| Hawai'i | Pu'u Maka'ala | 9 | 59 | RGCC001A_S25_ATGAG | A16 | AJR1017 |
| Hawai'i | Pu'u Maka'ala | 10 | NA | RGCC001A_S25_CAACC | A6 | AJR950 |
| Hawai'i | Pu'u Maka'ala | 11 | 85 | RGCC001A_S25_CGATC | A3 | AJR924 |
| Hawai'i | Pu'u Maka'ala | 12 | 80 | RGCC001A_S25_CTGCG | A17 | AJR1033 |
| Hawai'i | Pu'u Maka'ala | 13 | 60 | RGCC001A_S25_CTGTC | A18 | AJR1034 |
| Hawai'i | Pu'u Maka'ala | 14 | 52 | RGCC001A_S25_CTTGG | A19 | AJR1036 |
| Hawai'i | Pu'u Maka'ala | 15 | 74 | RGCC001A_S25_GACAC | A20 | AJR1037 |
| Hawai'i | Pu'u Maka'ala | 16 | NA | RGCC001A_S25_GCATG | A1 | AJR922 |
| Hawai'i | Pu'u Maka'ala | 17 | NA | RGCC001A_S25_GGTTG | A7 | AJR951 |
| Hawai'i | Pu'u Maka'ala | 18 | NA | RGCC001A_S25_TCGAT | A4 | AJR946 |
| Hawai'i | Pu'u Maka'ala | 19 | 61 | RGCC001A_S25_TGCAT | A5 | AJR947 |
| Hawai'i | Alili | 20 | 66 | RGCC001B_S26_AACCA | B2 | AJR839 |
| Hawai'i | Alili | 21 | 62 | RGCC001B_S26_AAGGA | B8 | AJR850 |
| Hawai'i | Alili | 22 | 88 | RGCC001B_S26_AATTA | B11 | AJR855 |
| Hawai'i | Alili | 23 | 100 | RGCC001B_S26_ACACA | B10 | AJR854 |
| Hawai'i | Alili | 24 | 45 | RGCC001B_S26_ACGGT | B12 | AJR857 |
| Hawai'i | Alili | 25 | NA | RGCC001B_S26_ACTTC | B14 | AJR859 |
| Hawai'i | Alili | 26 | 78 | RGCC001B_S26_AGCTA | B9 | AJR852 |
| Hawai'i | Alili | 27 | 118 | RGCC001B_S26_ATACG | B15 | AJR860 |
| Hawai'i | Alili | 28 | 104 | RGCC001B_S26_ATGAG | B16 | AJR861 |
| Hawai'i | Alili | 29 | 98 | RGCC001B_S26_CAACC | B6 | AJR848 |
| Hawai'i | Alili | 30 | NA | RGCC001B_S26_CGATC | B3 | AJR844 |
| Hawai'i | Alili | 31 | 107 | RGCC001B_S26_CTGCG | B17 | AJR891 |
| Hawai'i | Alili | 32 | NA | RGCC001B_S26_CTGTC | B18 | AJR893 |
| Hawai'i | Alili | 33 | NA | RGCC001B_S26_GACAC | B20 | AJR897 |
| Hawai'i | Alili | 34 | NA | RGCC001B_S26_GCATG | B1 | AJR838 |
| Hawai'i | Alili | 35 | 102 | RGCC001B_S26_GGTTG | B7 | AJR849 |
| Hawai'i | Alili | 36 | NA | RGCC001B_S26_TCGAT | B4 | AJR846 |
| Hawai'i | Alili | 37 | NA | RGCC001B_S26_TGCAT | B5 | AJR847 |
| Hawai'i | Kohala | 38 | NA | RGCC001C_S27_AACCA | C2 | AJR1413 |
| Hawai'i | Kohala | 39 | 55 | RGCC001C_S27_AAGGA | C8 | AJR1425 |
| Hawai'i | Kohala | 40 | 120 | RGCC001C_S27_AATTA | C11 | AJR1428 |
| Hawai'i | Kohala | 41 | 99 | RGCC001C_S27_ACACA | C10 | AJR1427 |
| Hawai'i | Kohala | 42 | 84 | RGCC001C_S27_ACTGG | C13 | AJR1430 |
| Hawai'i | Kohala | 43 | NA | RGCC001C_S27_ACTTC | C14 | AJR1476 |
| Hawai'i | Kohala | 44 | NA | RGCC001C_S27_AGCTA | C9 | AJR1426 |
| Hawai'i | Kohala | 45 | NA | RGCC001C_S27_ATACG | C15 | AJR1477 |
| Hawai'i | Kohala | 46 | NA | RGCC001C_S27_ATGAG | C16 | AJR1478 |
| Hawai'i | Kohala | 47 | NA | RGCC001C_S27_CAACC | C6 | AJR1417 |
| Hawai'i | Kohala | 48 | NA | RGCC001C_S27_CGATC | C3 | AJR1414 |
| Hawai'i | Kohala | 49 | NA | RGCC001C_S27_CTGCG | C17 | AJR1518 |
| Hawai'i | Kohala | 50 | NA | RGCC001C_S27_CTGTC | C18 | AJR1519 |
| Hawai'i | Kohala | 51 | NA | RGCC001C_S27_CTTGG | C19 | AJR1520 |
| Hawai'i | Kohala | 52 | NA | RGCC001C_S27_GCATG | C1 | AJR1412 |
| Hawai'i | Kohala | 53 | NA | RGCC001C_S27_GGTTG | C7 | AJR1418 |
| Hawai'i | Kohala | 54 | 117 | RGCC001C_S27_TCGAT | C4 | AJR1415 |
| Hawai'i | Kohala | 55 | NA | RGCC001C_S27_TGCAT | C5 | AJR1416 |
| Hawai'i | Ola'a | 56 | NA | RGCC001D_S28_AACCA | D2 | AJR1151 |
| Hawai'i | Ola'a | 57 | NA | RGCC001D_S28_AAGGA | D8 | AJR1158 |
| Hawai'i | Ola'a | 58 | 67 | RGCC001D_S28_AATTA | D11 | AJR1161 |
| Hawai'i | Ola'a | 59 | 64 | RGCC001D_S28_ACACA | D10 | AJR1160 |
| Hawai'i | Ola'a | 60 | 81 | RGCC001D_S28_ACTGG | D13 | AJR1187 |
| Hawai'i | Ola'a | 61 | 49 | RGCC001D_S28_ACTTC | D14 | AJR1188 |
| Hawai'i | Ola'a | 62 | 58 | RGCC001D_S28_AGCTA | D9 | AJR1159 |
| Hawai'i | Ola'a | 63 | 48 | RGCC001D_S28_ATACG | D15 | AJR1190 |
| Hawai'i | Ola'a | 64 | 72 | RGCC001D_S28_CAACC | D6 | AJR1156 |
| Hawai'i | Ola'a | 65 | 53 | RGCC001D_S28_CGATC | D3 | AJR1152 |
| Hawai'i | Ola'a | 66 | 119 | RGCC001D_S28_CTGCG | D17 | AJR1192 |
| Hawai'i | Ola'a | 67 | NA | RGCC001D_S28_CTGTC | D18 | AJR1232 |
| Hawai'i | Ola'a | 68 | NA | RGCC001D_S28_CTTGG | D19 | AJR1233 |
| Hawai'i | Ola'a | 69 | NA | RGCC001D_S28_GCATG | D1 | AJR1150 |
| Hawai'i | Ola'a | 70 | NA | RGCC001D_S28_GGTTG | D7 | AJR1157 |
| Hawai'i | Ola'a | 71 | NA | RGCC001D_S28_TCGAT | D4 | AJR1153 |
| Hawai'i | Ola'a | 72 | 109 | RGCC001D_S28_TGCAT | D5 | AJR1155 |
| Hawai'i | Thurston | 105 | NA | RGCC001H_S31_AACCA | H2 | AJR1061 |
| Hawai'i | Thurston | 106 | 44 | RGCC001H_S31_AAGGA | H8 | AJR1067 |
| Hawai'i | Thurston | 107 | 111 | RGCC001H_S31_AATTA | H11 | AJR1107 |
| Hawai'i | Thurston | 108 | 87 | RGCC001H_S31_ACACA | H10 | AJR1106 |
| Hawai'i | Thurston | 109 | NA | RGCC001H_S31_ACGGT | H12 | AJR1108 |
| Hawai'i | Thurston | 110 | NA | RGCC001H_S31_ACTGG | H13 | AJR1109 |
| Hawai'i | Thurston | 111 | NA | RGCC001H_S31_ACTTC | H14 | AJR1110 |
| Hawai'i | Thurston | 112 | NA | RGCC001H_S31_AGCTA | H9 | AJR1068 |
| Hawai'i | Thurston | 114 | 108 | RGCC001H_S31_CGATC | H3 | AJR1062 |
| Hawai'i | Thurston | 115 | 115 | RGCC001H_S31_GCATG | H1 | AJR1060 |
| Hawai'i | Thurston | 116 | 79 | RGCC001H_S31_GGTTG | H7 | AJR1066 |
| Hawai'i | Thurston | 117 | 97 | RGCC001H_S31_TGCAT | H5 | AJR1064 |
| Hawai'i | Saddle | 118 | NA | RGCC001I_S32_AACCA | H2 | AJRA2 |
| Hawai'i | Saddle | 119 | NA | RGCC001I_S32_CAACC | H6 | AJRA6 |
| Hawai'i | Saddle | 120 | 63 | RGCC001I_S32_CGATC | H3 | AJRA3 |
| Hawai'i | Saddle | 121 | 103 | RGCC001I_S32_GCATG | H1 | AJRA1 |
| Hawai'i | Saddle | 122 | NA | RGCC001I_S32_TCGAT | H4 | AJRA4 |
| Hawai'i | Saddle | 123 | 57 | RGCC001I_S32_TGCAT | H5 | AJRA5 |
| Hawai'i | Thurston | NA | 42 | RGCC001H_S31_TCGAT | H4 | AJR1063 |
| Hawai'i | Kohala | NA | 51 | RGCC001C_S27_ACGGT | C12 | AJR1429 |
| Hawai'i | Thurston | NA | 69 | RGCC001H_S31_CAACC | H6 | AJR1065 |
| Hawai'i | Alili | NA | 71 | RGCC001B_S26_ACTGG | B13 | AJR858 |
| Hawai'i | Alili | NA | 77 | RGCC001B_S26_CTTGG | B19 | AJR894 |
| Hawai'i | Pu'u Maka'ala | NA | 101 | RGCC001A_S25_AATTA | A11 | AJR949 |
| Hawai'i | Ola'a | NA | 113 | RGCC001D_S28_ACGGT | D12 | AJR1183 |
| Hawai'i | Kohala | NA | NA | RGCC001C_S27_GACAC | C20 | AJR1521 |
| Hawai'i | Ola'a | NA | NA | RGCC001D_S28_ATGAG | D16 | AJR1191 |
| Moloka'i | Moloka'i | 73 | NA | RGCC001F_S29_AACCA | F2 | RGGA10 |
| Moloka'i | Moloka'i | 74 | NA | RGCC001F_S29_AAGGA | F8 | RGG43 |
| Moloka'i | Moloka'i | 75 | 106 | RGCC001F_S29_AATTA | F11 | RGG46 |
| Moloka'i | Moloka'i | 76 | 86 | RGCC001F_S29_ACACA | F10 | RGG45 |
| Moloka'i | Moloka'i | 77 | 68 | RGCC001F_S29_ACGGT | F12 | RGG47 |
| Moloka'i | Moloka'i | 78 | 73 | RGCC001F_S29_ACTGG | F13 | RGG48 |
| Moloka'i | Moloka'i | 79 | NA | RGCC001F_S29_ACTTC | F14 | RGG5 |
| Moloka'i | Moloka'i | 80 | 114 | RGCC001F_S29_AGCTA | F9 | RGG44 |
| Moloka'i | Moloka'i | 81 | NA | RGCC001F_S29_ATACG | F15 | RGG6 |
| Moloka'i | Moloka'i | 82 | NA | RGCC001F_S29_ATGAG | F16 | RGG9 |
| Moloka'i | Moloka'i | 83 | 105 | RGCC001F_S29_CAACC | F6 | RGG41 |
| Moloka'i | Moloka'i | 84 | NA | RGCC001F_S29_CGATC | F3 | RGGA11 |
| Moloka'i | Moloka'i | 85 | NA | RGCC001F_S29_GCATG | F1 | RGGA9 |
| Moloka'i | Moloka'i | 86 | 112 | RGCC001F_S29_GGTTG | F7 | RGG42 |
| Moloka'i | Moloka'i | 87 | NA | RGCC001F_S29_TCGAT | F4 | RGG2 |
| Moloka'i | Moloka'i | 88 | NA | RGCC001F_S29_TGCAT | F5 | RGG3 |
| Maui | W_Maui | 89 | 56 | RGCC001G_S30_AACCA | G2 | RGG121 |
| Maui | W_Maui | 90 | NA | RGCC001G_S30_AAGGA | G8 | RGG135 |
| Maui | W_Maui | 91 | 43 | RGCC001G_S30_AATTA | G11 | RGG138 |
| Maui | W_Maui | 92 | 50 | RGCC001G_S30_ACACA | G10 | RGG137 |
| Maui | W_Maui | 93 | 116 | RGCC001G_S30_ACGGT | G12 | RGG151 |
| Maui | W_Maui | 94 | NA | RGCC001G_S30_ACTGG | G13 | RGG152 |
| Maui | W_Maui | 95 | 54 | RGCC001G_S30_ACTTC | G14 | RGG153 |
| Maui | W_Maui | 96 | 46 | RGCC001G_S30_AGCTA | G9 | RGG136 |
| Maui | W_Maui | 97 | 76 | RGCC001G_S30_ATACG | G15 | RGG154 |
| Maui | W_Maui | 98 | 41 | RGCC001G_S30_ATGAG | G16 | RGG155 |
| Maui | W_Maui | 99 | NA | RGCC001G_S30_CAACC | G6 | RGG133 |
| Maui | W_Maui | 100 | 110 | RGCC001G_S30_CGATC | G3 | RGG122 |
| Maui | W_Maui | 101 | NA | RGCC001G_S30_GCATG | G1 | RGGA2 |
| Maui | W_Maui | 102 | NA | RGCC001G_S30_GGTTG | G7 | RGG134 |
| Maui | W_Maui | 103 | NA | RGCC001G_S30_TCGAT | G4 | RGG131 |
| Maui | W_Maui | 104 | NA | RGCC001G_S30_TGCAT | G5 | RGG132 |
| Hawai'i | Ola'a | NA | NA | RGCC001D_S28_GACAC | D20 | AJR1186 |
| Hawai'i | Thurston | 113 | NA | RGCC001H_S31_ATACG | H15 | AJR1111 |

**Table S3:** F_ST_ values for ddRAD data from Ariamnes waikula spiders from Hawai’i.

|  | Alili | Kohala | Ola'a | Thurston | Saddle | Moloka'i | Maui |
| --- | --- | --- | --- | --- | --- | --- | --- |
| Pu'u Maka'ala | 0.0650 | 0.1466 | 0.0291 | 0.0461 | 0.1066 | 0.4870 | 0.5454 |
| Alili |  | 0.1394 | 0.0544 | 0.0629 | 0.0978 | 0.4829 | 0.5407 |
| Kohala |  |  | 0.1367 | 0.1496 | 0.1315 | 0.4742 | 0.5264 |
| Ola'a |  |  |  | 0.0334 | 0.0937 | 0.4817 | 0.5404 |
| Thurston |  |  |  |  | 0.1120 | 0.4815 | 0.5468 |
| Saddle |  |  |  |  |  | 0.4629 | 0.5410 |
| Moloka'i |  |  |  |  |  |  | 0.4781 |

**Table S4:** Average Tajima’s D values for each population of *A. waikula* (Hawai’i) and *A. melekalikimaka (*Maui) and *A. n. spp.* (Molokaʻi).

| Population | Average | Min | Max |
| --- | --- | --- | --- |
| Alili | -0.77 | -2.68 | 3.33 |
| Ola'a | -0.95 | -2.83 | 3.84 |
| Pu'u Maka'ala | -1.00 | -2.77 | 3.25 |
| Saddle | -0.61 | -2.97 | 3.79 |
| Thurston | -0.74 | -2.87 | 3.73 |
| Kohala | -0.71 | -2.82 | 3.52 |
| Maui | -0.06 | -2.63 | 3.94 |
| Moloka'i | -0.23 | -2.46 | 3.66 |

**Table S5:** Mean alpha diversity of the microbial communities per host populations. Alpha diversities were computed on rarefied OTU tables defined at 97% or Z-OTUs using the Chao 1 index.

| Island / species | Population | Number individuals | OTU at 97% |  | Z-OTU |  |
| --- | --- | --- | --- | --- | --- | --- |
|  |  |  | Alpha diversity | Standard deviation | Alpha diversity | Standard deviation |
| **Hawai'i**<br>*A. waikula* | **Alili** | 13 | 42.4 | 7.2 | 95.4 | 4.0 |
| **Hawai'i**<br>*A. waikula* | **Kohala** | 6 | 55.8 | 12.3 | 114.8 | 7.8 |
| **Hawai'i**<br>*A. waikula* | **Ola'a** | 9 | 41.6 | 3.0 | 96.3 | 8.5 |
| **Hawai'i**<br>*A. waikula* | **Pu'u Maka'ala** | 13 | 49.5 | 3.2 | 108.0 | 3.4 |
| **Hawai'i**<br>*A. waikula* | **Saddle** | 3 | 44.8 | 1.6 | 103.1 | 5.2 |
| **Hawai'i**<br>*A. waikula* | **Thurston** | 9 | 40.8 | 4.6 | 103.2 | 5.7 |
| **Maui**<br>*A. melekelikimaka* | **Puu Kukui** | 9 | 46.0 | 5.7 | 106.6 | 4.4 |
| **Moloka'i**<br>*A. n. sp.* | **Kamakou** | 7 | 52.2 | 6.7 | 106.6 | 5.3 |

**Table S6:** PERMANOVA testing the effect of host populations on the dissimilarities between microbial communities. The adjusted R-squared and the associated p-value are shown for each test. Significant relationships indicate a differentiation of the microbial communities according to the host populations.

| Populations | OTU at 97% |  |  | Z-OTU |  |  |
| --- | --- | --- | --- | --- | --- | --- |
|  | Whole microbiota | Gut microbiota | Endo-symbionts | Whole microbiota | Gut microbiota | Endo-symbionts |
| **All populations (from different species)** | 0.37905<br>(0.0001) | 0.31109<br>(0.0001) | 0.7146<br>(0.0001) | 0.37709<br>(0.0001) | 0.31739<br>(0.0001) | 0.7115<br>(0.0001) |
| ***A. waikula* populations within Hawai'i Island** | 0.21715<br>(0.0062) | 0.29671<br>(0.0007) | 0.28093<br>(0.0190) | 0.21542<br>(0.0057) | 0.29<br>(0.0003) | 0.29072<br>(0.0120) |

**Table S7:** Mantel tests comparing the microbial beta diversity dissimilarities to the genetic distances, the phylogenetic distances of their associated *Ariamnes,* and the geographical distances between the sampling sites. Beta diversity dissimilarities were computed on rarefied OTU tables defined at 97% using the Bray-Curtis dissimilarities. Bold values represent significant correlations. Mantel tests were either performed on all populations (between *Ariamnes* species and *A. waikula* populations) or only on *A. waikula* populations.

| Matrix 1 | Matrix 2 | All populations |  | *A. waikula* populations |  |
| --- | --- | --- | --- | --- | --- |
|  |  | Correlation | p-value | Correlation | p-value |
| Whole microbiota<br>beta diversity<br>distances | *Ariamnes*<br>genetic<br>distances | 0.312 | 9.9e-05 | 0.12 | 0.07 |
|  | *Ariamnes*<br>phylogenetic<br>distances | 0.323 | 9.9e-05 | 0.03 | 0.32 |
|  | Geographic<br>distances | 0.292 | 9.9e-05 | 0.06 | 0.16 |
| Endosymbionts<br>beta diversity<br>distances | *Ariamnes*<br>genetic<br>distances | 0.620 | 9.9e-05 | 0.01 | 0.50 |
|  | *Ariamnes*<br>phylogenetic<br>distances | 0.646 | 9.9e-05 | 0.06 | 0.26 |
|  | Geographic<br>distances | 0.565 | 9.9e-05 | 0.12 | 0.14 |
| Gut microbiota<br>beta diversity<br>distances | *Ariamnes*<br>genetic<br>distances | -0.096 | 0.84 | -0.04 | 0.67 |
|  | *Ariamnes*<br>phylogenetic<br>distances | -0.095 | 0.86 | -0.13 | 0.91 |
|  | Geographic<br>distances | -0.030 | 0.62 | -0.07 | 0.73 |

